# Structural Remodeling of Fungal Cell Wall Promotes Resistance to Echinocandins

**DOI:** 10.1101/2023.08.09.552708

**Authors:** Malitha C. Dickwella Widanage, Isha Gautam, Daipayan Sarkar, Frederic Mentink-Vigier, Josh V. Vermaas, Thierry Fontaine, Jean-Paul Latgé, Ping Wang, Tuo Wang

## Abstract

The insufficient efficacy of existing antifungal drugs and the rise in resistance necessitate the development of new therapeutic agents with novel functional mechanisms^1,2^. Echinocandins are an important class of antifungals that inhibit β-1,3-glucan biosynthesis to interfere with cell wall structure and function^3,4^. However, their efficacy is limited by the fungistatic activity against *Aspergillus* species and the trailing effect during clinical application. Here, we describe how echinocandins remodel the supramolecular assembly of carbohydrate polymers in the fungal cell wall in an unexpected manner, possibly resulting in a subsequent inhibition of the activity of these drugs. Solid-state nuclear magnetic resonance (ssNMR) analysis of intact cells from the human pathogenic fungus *Aspergillus fumigatus* showed that the loss of β-1,3-glucan and the increase of chitin content led to a decrease in cell wall mobility and water-permeability, thus enhancing resistance to environmental stresses. Chitosan and α-1,3-glucan were found to be important buffering molecules whose physical association with chitin maintained the wall integrity. These new findings revealed the difficult-to-understand structural principles governing fungal pathogens’ response to echinocandins and opened new avenues for designing novel antifungal agents with improved efficacy.

---

Invasive fungal infections have high occurrence and mortality among immunocompromised patients^5,6^. SARS-CoV-2–associated fungal infections have significantly impacted a substantial portion of critically ill individuals suffering from COVID-19^7,8^. There are currently a few antifungal agents available, with often therapeutic failures and a recently observed increase in drug resistance^9,10^. Azoles, the primary antifungal drugs that target the ergosterol biosynthesis in fungi, are facing rising resistance, underscoring the need for novel therapeutic agents^11,12^. Consequently, echinocandins, including caspofungin, micafungin, and anidulafungin, have been approved for clinical applications^13^. These cyclic hexapeptides possess modifiable aliphatic tails (**Fig. 1a**) and inhibit the biosynthesis of β-1,3-glucan, a major and widespread structural polysaccharide in the fungal cell wall^3,14^. The unique target has granted these lipopeptides remarkable efficacy and safety. As an effective fungicide for *Candida* species, echinocandins are being used as preferred agents for treating invasive candidiasis^15,16^. However, echinocandins only have relatively low efficacy when treating *A. fumigatus*, the most ubiquitous airborne fungal pathogen responsible for pulmonary infections and invasive aspergillosis^17,18^. Such discrepancy should result from the distinct composition and organization of the cell walls in *Candida* and *Aspergillus* species.

**Figure 1.**
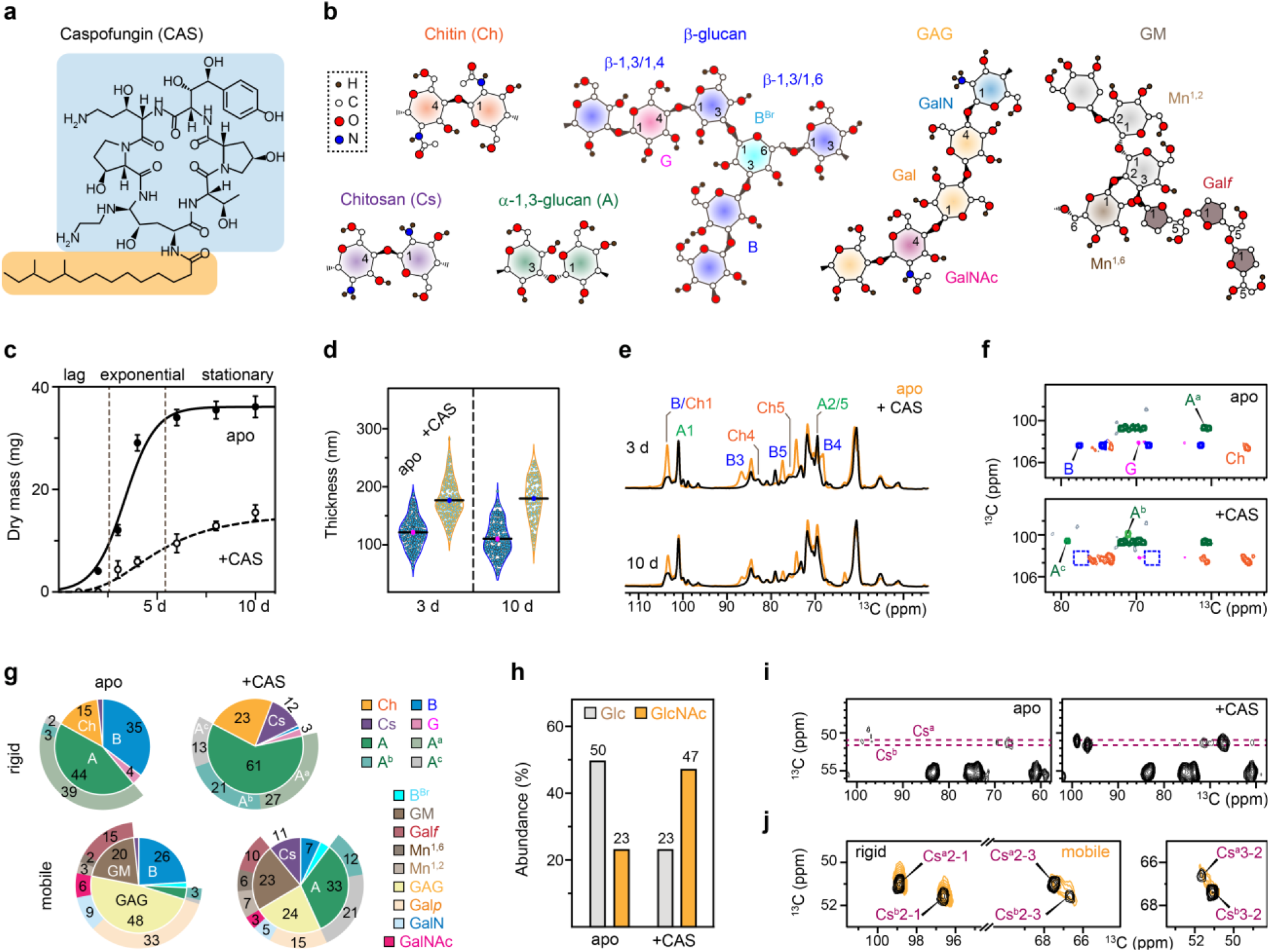
Alternation of cell wall polymer composition due to caspofungin treatment. **a**, Chemical structure of caspofungin (CAS), highlighting the cyclic peptide in blue and the lipid component in yellow. **b**, Simplified structures of fungal cell wall polysaccharides. NMR abbreviations are provided for each polysaccharide or monosaccharide unit. **c**, Growth profiles of *A. fumigatus* without (apo) and with caspofungin as a change in dry mass. **d**, Inner cell wall thickness determined using TEM images. **e**, 1D ^13^C CP spectra of four *A. fumigatus* samples showing the compositional difference of rigid polysaccharides. **f**, Comparison of 2D ^13^C-^13^C spectra of different cell walls. The missing peaks of β-1,3-glucans in drug-treated samples are highlighted using blue boxes. **g**, Molar composition of rigid (top) and mobile (bottom) cell wall polysaccharides estimated using volumes of resolved cross peaks in 2D ^13^C CORD and DP refocused J-INADEQUATE spectra. Ch: chitin, Cs: chitosan, B: β-1,3-glucan, G: β-1,4-glucose residue, A: α-1,3-glucan. For α-1,3-glucan in the rigid portion, the inner pie chart displays the total amount, while the outer circle shows the individual content of three subtypes (A^a^, A^b^, and A^c^). B^Br^: β-1,3,6-glucose residue (the branching point), GM: galactomannan, Gal*f*: galactofuranose, Mn^1,2^: α-1,2-mannose, Mn^1,6^: α-1,6-mannose. For the mobile phase, the inner pie chart depicts the total content of each polysaccharide, while the outer circle shows the monosaccharide units or subtypes. **h**, Changes of the total Glc and GlcNAc in 3-day-old cell wall using GC-MS. **i**, Detection of two chitosan types (Cs^a^ and Cs^b^) in drug-treated 3-day-old sample in 2D CORD spectra. **j**, Presence of both chitosan forms in 3-day-old caspofungin-treated sample across both rigid fraction (black; CORD spectra) and mobile fraction (yellow; sheared refocused DP J-INADEQUATE spectra). For **c** and **d**, data are mean ± s.e. *n* = 3 replicates for **c** and *n* = 200 in 10 cells for each sample in **d**. Source data are provided as a Source Data file.

Alongside β-1,3-glucans, the major carbohydrate components of *A. fumigatus* encompass α-1,3-glucan, chitin, galactomannan (GM), and galactosaminogalactan (GAG) (**Fig. 1b**)^19^. Although with a lower occurrence, β-1,6- and 1,4-linkages are also present in glucans, with the former forming the branching anchor in β-1,3/1,6-glucans and the latter existing in the β-1,3/1,4-glucan domain or serving as a linker to other polysaccharides, such as chitin^20^. Echinocandins only exert a fungistatic effect on *A. fumigatus*, causing lysis to the hyphal tips without completely preventing growth^21^. This phenomenon has been postulated to stem from the perturbations of the membrane environment that encloses glucan synthases or from the restructuring of the cell wall, but there is no molecular-level evidence supporting these hypotheses^22–24^.

Recently, magic-angle spinning (MAS) ssNMR methods have been established to uncover the cell wall architecture in *Aspergillus* and other fungal species^25–27^. High-resolution data collected on intact and uniformly ^13^C,^15^N-labeled *A. fumigatus* mycelia have demonstrated that α-1,3-glucan associates with chitin to form a robust mechanical scaffold, which is dispersed within a mobile and hydrated mesh formed by branched β-glucans, and enclosed by an dynamic outer shell rich in GM and GAG^28,29^. This nanoscale architecture undergoes significant rearrangements in response to biosynthesis deficiencies in structural polysaccharides and during morphotype transitions throughout fungal life cycles^30,31^. Leveraging these advancements, our objective here is to investigate a question of both biochemical and medical significance: How do fungal pathogens restructure their cell walls to develop resistance against antifungal drugs?

SsNMR and dynamic nuclear polarization (DNP) techniques, in conjunction with transmission electron microscope (TEM) imaging, biochemical analysis, and molecular dynamics (MD) simulations, have unveiled that the echinocandin-induced augmentation of the chitin content results in enhanced stiffness and hydrophobicity of the polymer network within the cell wall. The rise in chitin content is associated with the deacetylation of these polysaccharides, leading to a significant amount of chitosan in the treated fungus. The physical associations of chitosan, α-1,3-glucan, and chitin provide critical mechanical support, essential for preserving cell wall integrity. While only a low amount of β-glucan remains, it exhibits an atypical structure characterized by a high degree of branching. To compensate for the loss of most β-glucans, two novel types of semi-dynamic α-1,3-glucans are produced to regenerate the soft matrix. Additionally, reshuffling the composition and structure of GAG and GM occurs on the cell surface. These modifications by echinocandin treatment provide heretofore unavailable molecular insights into the limited effectiveness of existing antifungal drugs and establish the structural foundation for developing novel antifungal compounds targeting the restructured polysaccharides and cell walls.

## Caspofungin alters the carbohydrate core

The fungal cell wall has a highly dynamic structure that undergoes constant modifications in response to external stresses^32–35^. The addition of 2.5 µg/mL caspofungin (above the minimum inhibitory concentration)^36,37^ suppressed the growth of *A. fumigatus* (**Fig. 1c**) and increased the thickness of the cell wall from 123-133 nm to 182-185 nm (**Fig. 1d** and **Supplementary Table 1**). Uniformly ^13^C, ^15^N-labeled *A. fumigatus* mycelia were therefore subjected to ssNMR analysis to identify molecular-level factors driving this microscopic-scale restructuring. Since caspofungin inhibits the biosynthesis of β-1,3-glucans in *A. fumigatus*, we particularly focused on the relatively rigid inner domains of cell walls, where β-1,3-glucans coexist with chitin and other glucans. Initial screening by one-dimensional (1D) ^13^C spectra confirmed the reduction of the β-1,3-glucan content. This decline was evident from the reduced intensity of characteristic peaks associated with β-1,3-glucan carbon 3 (B3) at 86 ppm, carbon 4 (B4) at 69 ppm, and carbon 5 (B5) at 77 ppm (**Fig. 1e**). For instance, the intensity of the B3 peak dropped by 94% in the echinocandin-treated sample. These changes were reproducibly observed across three sample batches (**Supplementary Fig. 1**).

Further two-dimensional (2D) ^13^C-^13^C/^15^N correlation spectra were acquired to resolve a large number of carbon sites in fungal glycans (**Extended Data Fig. 1a**). As anticipated, the signals of β-1,3-glucans became almost negligible in the drug-treated cell walls (**Fig. 1f**). Analysis of peak volumes revealed that caspofungin reduced the content of β-1,3-glucan in the rigid portion of the cell wall from 35% to 1% in young (3 day) cultures, and from 16% to 1% in older (10 day) cultures (**Fig. 1g** and **Extended Data Fig. 1b**). In contrast, the quantity of chitin increased by 1.5-2-fold in these cultures (**Fig. 1g** and **Supplementary Table 2**). This is not surprising because the increase of chitin is a classical compensatory reaction for *A. fumigatus* to respond to many stresses^4,37,38^. Chemical analysis of the sugar composition through GC-MS and HPLC chromatography validated the reduction in β-glucan content and the elevation of chitin level in the cell wall (**Fig. 1h** and Supplementary Table 3).

Interestingly, the content of chitosan, the deacetylated form of chitin, was initally minimal in the apo sample, but after caspofungin treatment, it increased to comprise 11-12% of the cell wall carbohydrates (Fig. 1i). Two distinct forms of chitosan molecules were identified, equally populated and evenly distributed across both the rigid and mobile fractions of the cell wall (Fig. 1j). Although the structural role of chitosan in *Aspergillus* species has long been overlooked, emerging data is shedding light on its association with external stressors, including hypersaline conditions, as reported recently^39^, and caspofungin exposure, as demonstrated here.

The atomic resolution of ssNMR offers a novel capability to discern intricate structural characteristics of β-glucan complex within intact *A. fumigatus* cell walls (Fig. 2a). This includes spectroscopic identification of glucopyranose residues bearing β-1,3, β-1,3,6, and β-1,4 linkages, with a particular focus on distinguishing the branching points (B^Br^) from the predominant linear chains (Fig. 2b and **Extended Data Fig. 1c**). The residual β-glucan surviving caspofungin treatment was hyperbranched through many β-1,3,6-linkages while its content of β-1,3/1,4-linked sequence was lower (Fig. 2c and **Supplementary Fig. 2**). This finding suggests that β-1,3/1,4-glucan polymerase, coded by the gene TFT1^40^, was sensitive to caspofungin, or its expression was downregulated in the presence of caspofungin. The increased branching of β-1,3-glucan as a caspofungin-remodeled structure suggests that extracellular remodeling process, which requires members of the glycoside hydrolase GH-72 family^41^, remain unaltered. Cross-linking of GM onto the cell wall β-1,3-glucan represents another crucial remodeling process that occurs at the extracellular level^42^. In caspofungin-treated mycelia, the ratio between GM and β-1,3-glucan was elevated (Fig. 1g), suggesting that the high level of β-1,3-glucan branching and the GM-β-1,3-glucan reticulation could act as compensatory effects following the inhibition of β-1,3-glucan synthesis to support the vegetative growth of *A. fumigatus* under caspofungin treatment.

**Figure 2.**
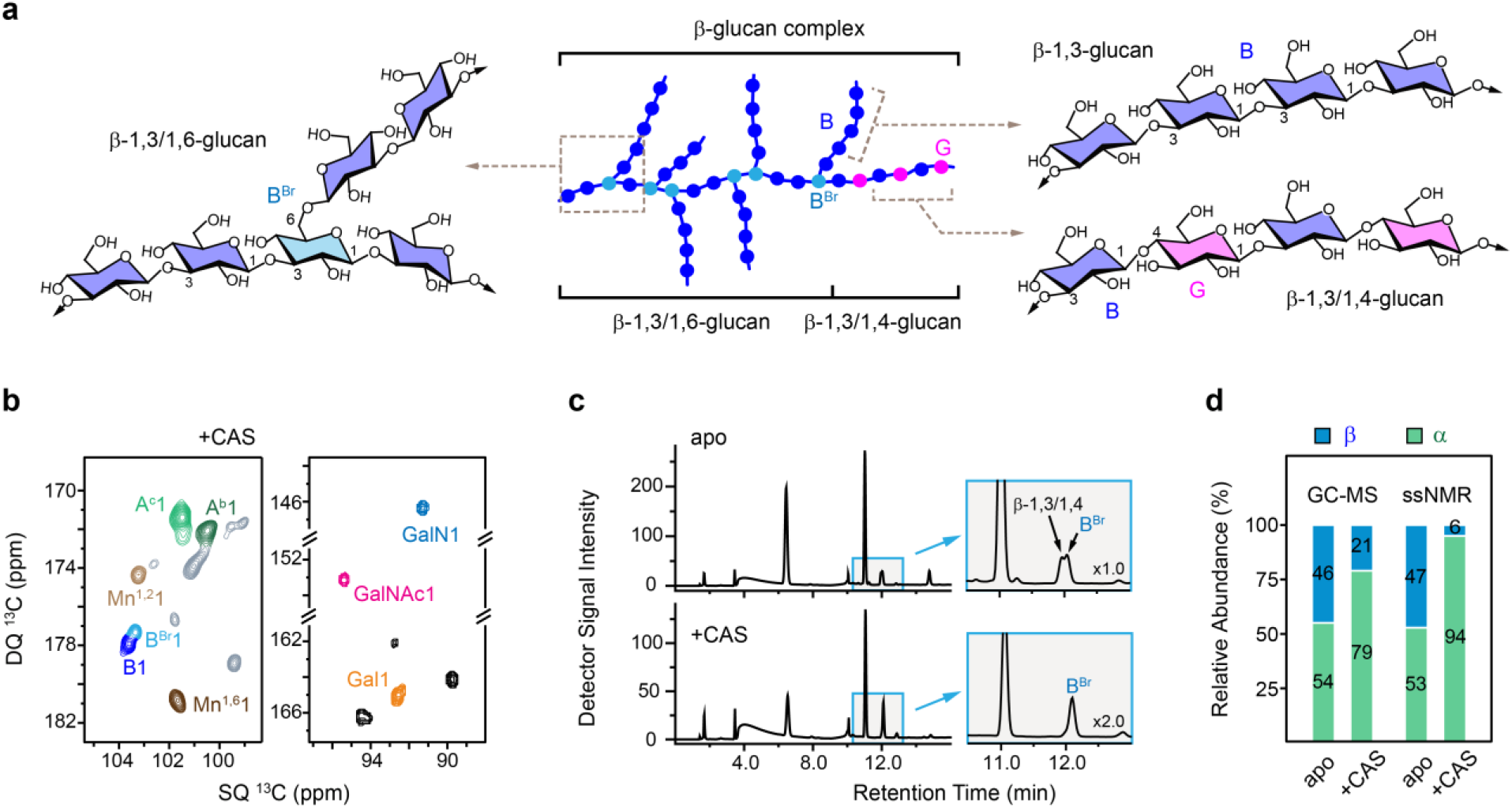
Caspofungin remodels the structures of β- and α-glucans. **a**, Diagram illustrating the complex structure of β-glucans in *A. fumigatus* cell walls. NMR abbreviations are introduced to annotate different linkages in the main chains and branches. **b**, Mobile polysaccharides detected using 2D ^13^C DP J-INADEQUATE spectrum resolving the β-1,3-linked (B) and β-1,3,6-linked (B^Br^) glucose units, as well as two novel forms of α-1,3-glucans (A^a^ and A^b^). **c**, Qualitative analysis by HPLC shows the presence of β-glucan complex and the reduced amount of β-1,3/1,4-glucan domains after caspofungin treatment in 3-day-old cultures. **d**, SsNMR and GC-MS analysis of 3-day-old cell walls, with the results expressed as percentages of the total cell wall glucans.

Caspofungin dramatically increased the amount of α-1,3-glucan in 3-day-old samples, from 44% to 61% and from 4% to 33% in the rigid and mobile fractions, respectively (Fig. 1g and **Supplementary Table 4**). The accumulation of α-glucan compared to β-glucans was further validated using chemical assays (Fig. 2d and **Supplementary Table 5**). Apart from the major type of α-1,3-glucan (type-a or A^a^), two other magnetically inequivalent forms (A^b^ and A^c^) were observed in the caspofungin-treated sample, which was present in both the rigid domain (Fig. 1f) and the mobile fraction (Fig. 2b). These two new types of α-1,3-glucans have not been reported previously and their chemical identities were confirmed by their absence from an *A. fumigatus* mutant and other fungal species, such as *Candida albicans* and *Aspergillus sydowii*^39^, which lack α-1,3-glucan (**Extended Data Fig. 2**). The three types of α-1,3-glucans exhibited distinct C3 chemical shift, ranging from 85 ppm for type-a to 79-81 ppm for types-b and c. Since C3 denotes the carbon position of the glycosidic linkage in this polysaccharide, these observed differences imply alternations in the helical screw conformation. Analogous conformational variations have been noted in plant polysaccharides such as xylan and homogalacturonan due to binding with other polysaccharides or diatomic cations^43–45^. While the type-b and c forms exhibited very weak signals in the apo sample, they showed strong peaks in the *A. fumigatus* cell walls after caspofungin treatment (**Extended Data Fig. 3**). Thus, α-1,3-glucan simultaneously increased its amount and altered its structure to compensate for the loss of β-1,3-glucan.

Other mobile polysaccharides, primarily GAG and GM, were largely conserved as shown by their comparable chemical shift fingerprints (**Supplementary Fig. 3**). Caspofungin halved the GAG content but did not perturb the molecular composition of the three sugar residues (Gal*p*, GalN, and GalNAc) within this heteroglycan (Fig. 1g and **Supplementary Table 3**). Due to the cationic state of GalN (present as GalNH_3_^+^) at cellular pH, the observed decrease in the GAG content will reduce the charge of the cell wall surface and affect the adhesive property. Meanwhile, GM now expects shorter and fewer sidechains, indicated by the reduced amount of Gal*f* (Fig. 1g). The ratio of carbohydrates to proteins/lipids remained consistent in the rigid portions of all four samples, indicating that the total amount of structural polysaccharides was effectively maintained even after caspofungin exposure (**Supplementary Fig. 4** and **Table 6**).

## Reduced water retention and polymer dynamics

A key finding is that caspofungin treatment leads to a reduction in the water accessibility of *A. fumigatus* cell walls, which was marked by the decline of intensities in 2D hydration maps (Fig. 3a). Such intensities reflect the capabilities of polymers to retain water molecules around various carbon sites (**Supplementary Table 7**). Upon caspofungin treatment, the relative intensities of α-glucan, chitin, and β-1,3-glucan in cell walls of 3-day-old cultures dropped by 27%, 39%, and 46%, respectively (Fig. 3b). This decrease in water retention and increase in hydrophobicity should contribute to the reduction in cell wall permeability.

**Figure 3.**
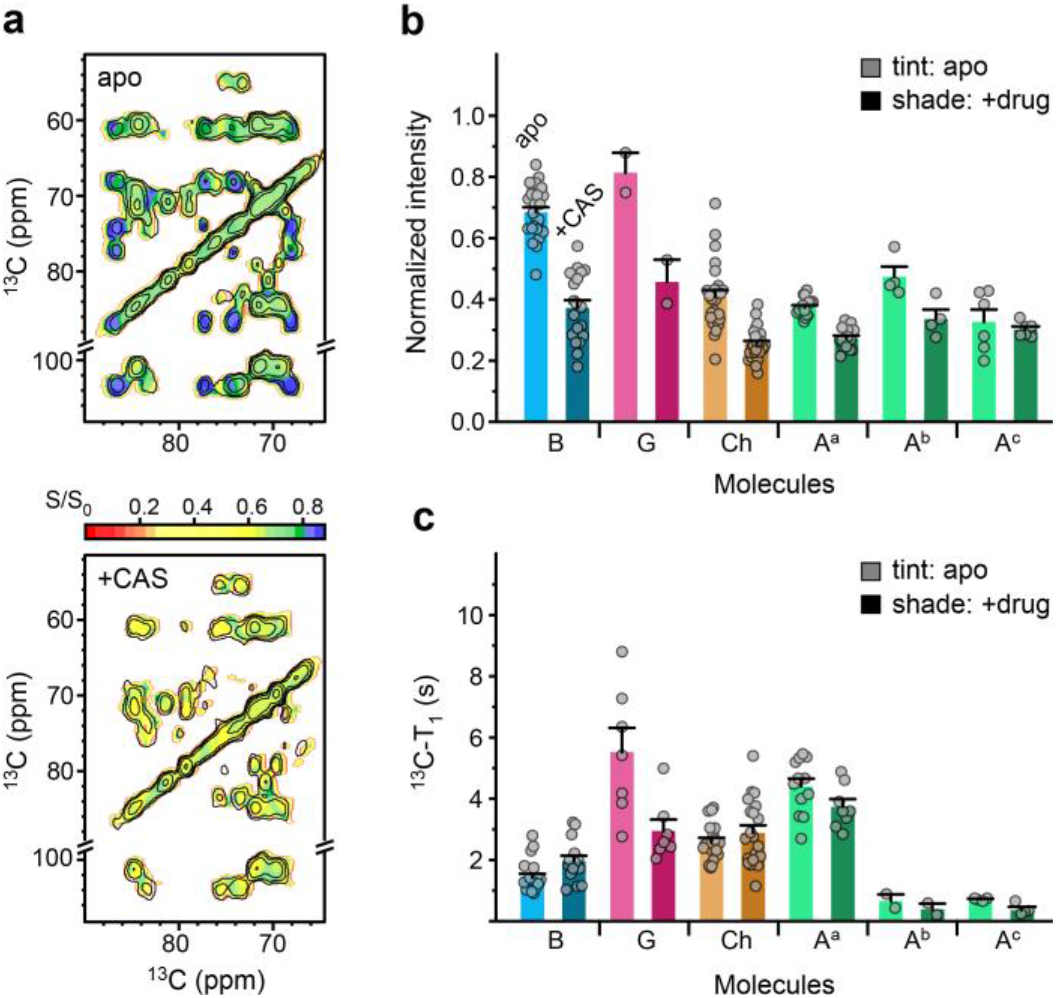
Regulation of water accessibility and biopolymer rigidity by caspofungin. **a**, Hydration map of 3-day-old *A. fumigatus* without (top) and with (bottom) caspofungin treatment. This intensity map plotted the ratios (S/S_0_) of peak intensities from the water-edited spectrum (S) detecting hydrated molecules, relative to those from the control spectrum (S_0_) representing equilibrium conditions. **b**, Relative intensities (S/S_0_) of different carbon sites indicating the extent of water association of cell wall polysaccharides. **c**, ^13^C-T_1_ relaxation time constants for different carbon sites in 3-day-old *A. fumigatus* samples with and without caspofungin treatment. Data are presented as mean ± s.e. and *n* varies for each component, with data points superimposed on bars. Source data are provided as a Source Data file.

The dynamical heterogeneity of cell wall polysaccharides was examined using ^13^C spin-lattice (T_1_) relaxation measurements, carried out in a pseudo-3D format to enhance spectral resolution (**Supplementary Fig. 5**). Within each sample, β-1,3-glucans showed high mobility on the nanosecond timescale, with relatively short ^13^C-T_1_ time constants of 1-3 s (Fig. 3c and **Supplementary Table 8**). Chitin, the major form of α-1,3-glucan (A^a^), and β-1,4-linked glucose units exhibited longer ^13^C-T_1_ values of 3-8 s. The stiffness of chitin and α-1,3-glucan is justified by their close association at the mechanical center, but the long T_1_ values of β-1,4-linked glucose units, typically identified in the mix-linked β-1,3/1,4-glucan, are unexpected^28^. These β-1,3/1,4-glucans constitute minor segments that are occasionally covalently connected to the major β-1,3/1,6-glucan domains^20^. The data suggest that β-1,3/1,4-glucan is intrinsically inflexible, either achieved by maintaining physical separation from the bulk domain of mobile β-glucans or due to spatial constraints imposed by packing interactions with rigid molecules like chitin or α-glucan.

Caspofungin reshapes the dynamical landscape of the mechanical core in *A. fumigatus* cell walls. After treatment, the residual β-1,3-glucan (which transforms into highly branched β-1,3/1,6-glucan) was rigidified, with its average ^13^C-T_1_ increased apparently from 1.4-s to 2.0-s (Fig. 3c). Because most β-1,3-glucans residing in the soft matrix were eliminated due to drug inhibition, the remaining ones should primarily pack with rigid molecules. Caspofungin also rigidified chitin and increased its ^13^C-T_1_ from 2.6-s to 2.9-s. This could be attributed to the promoted aggregation of chitin chains in the absence of filling molecules (β-1,3-glucans). Conversely, β-1,4-linked glucose residues experienced an opposite shift, with almost halved ^13^C-T_1_ (from 5.5-s to 3.0-s). The removal of β-1,3-glucans has significantly impacted the stability of β-1,3/1,4-glucans, probably because these two polysaccharides were covalently linked to each other^20,40^. Similarly, the major form of α-1,3-glucan became moderately more dynamic, and the two minor forms are inherently dynamic, which should help to partially fulfill the diverse roles of the now-removed β-1,3-glucan.

## Rearrangement of the macromolecular network

Fungal cell walls undergo significant spatial rearrangement in response to caspofungin. Assessment of the nanoscale assembly of macromolecules is enabled by the DNP technique, which relies on electron polarization to provide a 17-22-fold enhancement on the NMR sensitivity of two 3-day-old *A. fumigatus* samples (Fig. 4a)^46–48^. This enhancement results in a shortened experimental duration by 290 to 480 times. The cryogenic temperature required for DNP measurements has broadened the NMR linewidth and abolished the signals of highly disordered molecules such as GM and GAG, but the resolution remains adequate for resolving the diverse carbon and nitrogen sites of major polysaccharides and visualizing their sparsely populated packing interfaces (**Extended Data Fig. 4**).

**Figure 4.**
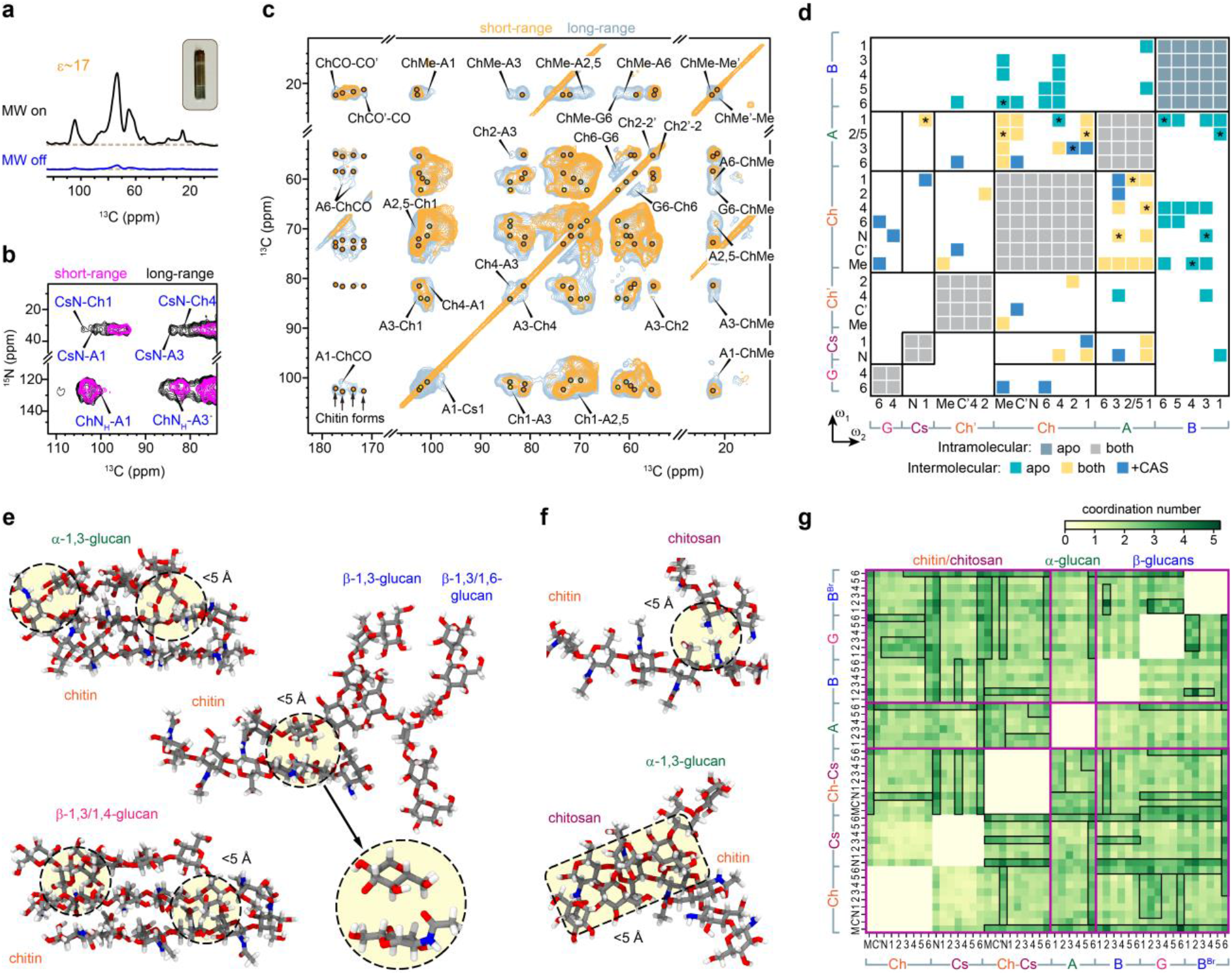
Intermolecular contacts in 3-day-old apo and caspofungin-treated *A. fumigatus*. **a**, DNP enhances NMR sensitivity of the caspofungin-treated 3-day-old sample by 17-fold when microwave (MW) is activated. Inset shows the DNP sample with 30 mg hydrated mycelial material enclosed in a 3.2-mm sapphire rotor. Dash lines mark the baseline of the spectra. **b**, DNP 2D ^15^N-^13^C correlation spectra of caspofungin-treated sample. Interactions happen between the ^15^N-site of the chitin amide (ChN_H_) or chitosan amine (CsN) and the carbons of polysaccharides. For example, CsN-A1 represents the cross peak between chitosan nitrogen with α-1,3-glucan carbon 1. **c**, DNP 2D ^13^C-^13^C correlation spectra of caspofungin-treated sample. Most interactions happen between α-glucan and chitin. The carbonyl region showed four types of chitin signals. **d**, Site-specific summary of intermolecular cross peaks identified among different polysaccharides. Diagonal regions exhibit intramolecular cross peaks. Off-diagonal regions show intermolecular interactions happening only in the apo sample (green), only in the drug-treated sample (blue), or in both samples (yellow). Strong intermolecular interactions from short-mixing spectra are marked with asterisks. The plot can be asymmetric relative to the diagonal due to the directionality of polarization transfer, e.g., Ch1-A3 differs from A3-Ch1. Left and bottom axes indicate the carbohydrate carbon numbers observed in indirect (ω_1_) and direct (ω_2_) spectral dimensions. Representative short-range interactions observed during 1 *μ*s all-atom MD are shown between **e**, chitin and glucans, and **f**, chitosan and other polymers. Atoms in gray, red, blue, and white represent carbon, oxygen, nitrogen, and hydrogen, respectively. Packing interactions within 5 Å are highlighted. **g**, Contact map of intermolecular interactions within 5 Å identified in MD models. Coordination number (see Methods) represents the number of contacts between two carbon sites of different polysaccharides. Magenta lines separate the sections of chitin/chitosan (Ch: chitin; Cs: chitosan; Ch-Cs: chitin-chitosan copolymer), α-1,3-glucan (A), and β-glucans (B: linear β-1,3-glucan; G: β-1,3/1,4-glucan; B^Br^: β-1,3/1,6-glucan). Carbon and nitrogen numbers are provided on both axes (M: Me or methyl; C’ carbonyl). Regions boxed in black highlight extensive interactions.

Caspofungin-treated samples showed unambiguous cross peaks between the nitrogen of chitosan amine group (CsN) and the carbons of chitin and α-glucan (Fig. 4b), providing clear evidence of the nanoscale coexistence of these biomolecules. Notably, the amide nitrogen of chitin correlates predominantly with the carbons of α-1,3-glucan, forming a physically associated complex. This structural concept is further supported by the numerous ^13^C-^13^C cross peaks between these two polysaccharides (Fig. 4c). In addition, four major forms of N-acetylglucosamine (GlcNAc) units were identified. These units engage in hydrogen-bonding to create crystalline microfibrils^49^, and inter-form cross peaks can be detected between the carbonyls (ChCO-CO’) and between the methyl groups (ChMe-Me’) of different conformers.

A statistical summary of 114 intermolecular cross peaks revealed the elimination of intermolecular contacts involving β-1,3-glucans, which is an expected outcome due to the depletion of this polysaccharide by caspofungin (Fig. 4d and **Extended Data Fig. 5**). In contrast, drug treatment promoted the extensive associations between chitin and α-1,3-glucans, thereby stabilizing the polymer complexes formed by these two macromolecules. Chitosan was found to be anchored to this chitin-α-1,3-glucan core, and caspofungin treatment has strengthened this packing interaction. Chitosan could potentially exist as a constituent of chitin-chitosan copolymers that are sometimes referred to as partially deacetylated chitin, or as fully deacetylated chains that are physically deposited onto the surface of chitin crystallites. Interestingly, β-1,4-linked glucose residues also showed several cross peaks with chitin. This suggests that β-1,3/1,4-glucan is trapped by chitin microfibrils, which explains the distinctive rigidity of this linear polymer (Fig. 3c). Therefore, in the absence of the bridging molecule β-1,3-glucan, the stability of *A. fumigatus* cell walls are upheld through many inter-allomorph interactions within chitin crystallites and intermolecular associations involving chitin, α-1,3-glucans, chitosan, and β-1,3/1,4-glucan, collectively ensuring the maintenance of structural integrity.

Despite the potential involvement of protein and lipid in the cell wall architecture, no observable interactions were detected between these non-carbohydrate components and the cell wall polysaccharides (**Supplementary Fig. 6**). Recent chemical and NMR analyses have reported covalent connections between structural proteins and GM/GAG through unusual alkali-resistant linkers that incorporate hydrophobic amino acid residues such as valines^31^. This structural pattern remains unaffected in the caspofungin-treated cell wall, as evident from the consistent signals of valine and other amino acids in the rigid phase of the *A. fumigatus* samples (**Supplementary Fig. 4d**). It can be inferred that lipids and proteins have limited contact with cell wall polysaccharides, primarily associating with GM and GAG in the surface layer.

Microsecond-long all-atom molecular dynamics (MD) simulations have been employed to quantify intermolecular interactions and visualize polysaccharide packing interfaces in cell walls^50,51^. The solvated molecular system includes 14 polysaccharides, with two copies of each linear polysaccharides (chitin, chitosan, chitin-chitosan copolymer, α-1,3-glucan, β-1,3-glucan, and β-1,3/1,4-glucan) and the branched polymer β-1,3/1,6-glucan (**Supplementary Movie 1** and Fig. 7). From microsecond-long unbiased MD trajectory, we systematically scanned over different cutoff distances to locate the best-fit trajectory, considering both short-range interactions (<7 Å) and long-range interactions (7-10 Å) based on the experimental observations provided by ssNMR. Chitin physically aggregated with both α-1,3-glucan and the linear terminal chain of β-1,3/1,4-glucan (Fig. 4e and **Supplementary Movie 2**). The intermolecular distances are typically within 5.0 Å in the molecular trajectories, which rationalizes the strong cross peaks observed among these polysaccharides in caspofungin-treated cell walls. Chitin also exhibited close range interactions with both linear β-1,3-glucan and branched β-1,3/1,6-glucan; however, these molecules tend to be dispersed in space (Fig. 4e and **Supplementary Movie 3**). Chitin’s association with chitosan was stable, with nitrogen-carbon/nitrogen distances of less than 5.0 Å (Fig. 4f and **Supplementary Movie 4**), highlighting the importance of hydrogen bonds involving nitrogen sites in chitin amide and chitosan amine. Short-range interactions were also observed between chitin-chitosan copolymers and α-1,3-glucans (Fig. 4f). Two chitin polymers also tend to exist as a stacked assembly (**Supplementary Movie 5**), which can serve as the platform to accommodate external molecules, such as α-1,3-glucan and β-1,3/1,4-glucan as shown in Fig. 4e. These simulation results explained the origin of NMR cross peaks happening between the chitosan introduced by caspofungin treatment and the stiff scaffold of chitin and α-glucan in the cell wall.

The interatomic contact map (Fig. 4g) allows us to visualize extensive short-range interactions (within 5 Å) between the chitin methyl group and the pyranose rings of both α-1,3- and β-glucans. The amine nitrogen of chitosan also played a major role in interacting with all three types of β-glucans. α-1,3-glucan is capable of interacting with both chitin and chitosan, utilizing its carbon-6 that extends outside the pyranose ring. Alternatively, all carbon sites of α-1,3-glucan predominantly interacted with the chitin-chitosan copolymer, a component that is possibly present only in the caspofungin-treated *A. fumigatus* cells. Therefore, the NMR-observed strong cross peaks between α-1,3-glucan and chitin/chitosan after the depletion of β-1,3-glucan by caspofungin were not surprising. These unbiased all-atom MD models provided detailed structural insights into the pairwise interactions between different functional groups present in fungal polysaccharides.

## Structural mechanism of echinocandin resistance

The fungal cell wall is an elastic entity that withstands hydrostatic turgor pressure, with microfibrillar components restricting stretching and matrix components countering compression^52^. Most polymers are interconnected by covalent linkages and held together by hydrogen bonds rather than as separate components^53^. Previous chemical and imaging studies and recent NMR data showed that the *A. fumigatus* mycelial cell wall comprises an outer shell primarily composed of GAG, GM, and associated proteins, and an inner layer consisting of a β-glucan network that encompasses the hydrophobic and rigid junctions formed by physically packed chitin and α-glucans (Fig. 5a)^29,31^. At the same time, β-1,3-glucan also covalently crosslinks chitin and GM together, as revealed by chemical analysis^19^. The current study further reshapes our perception of the response of fungal cell wall to antifungal treatment and reveals that *A. fumigatus* relies on the development of a thicker, stiffer, and more dehydrated cell wall to counter external stresses.

**Figure 5.**
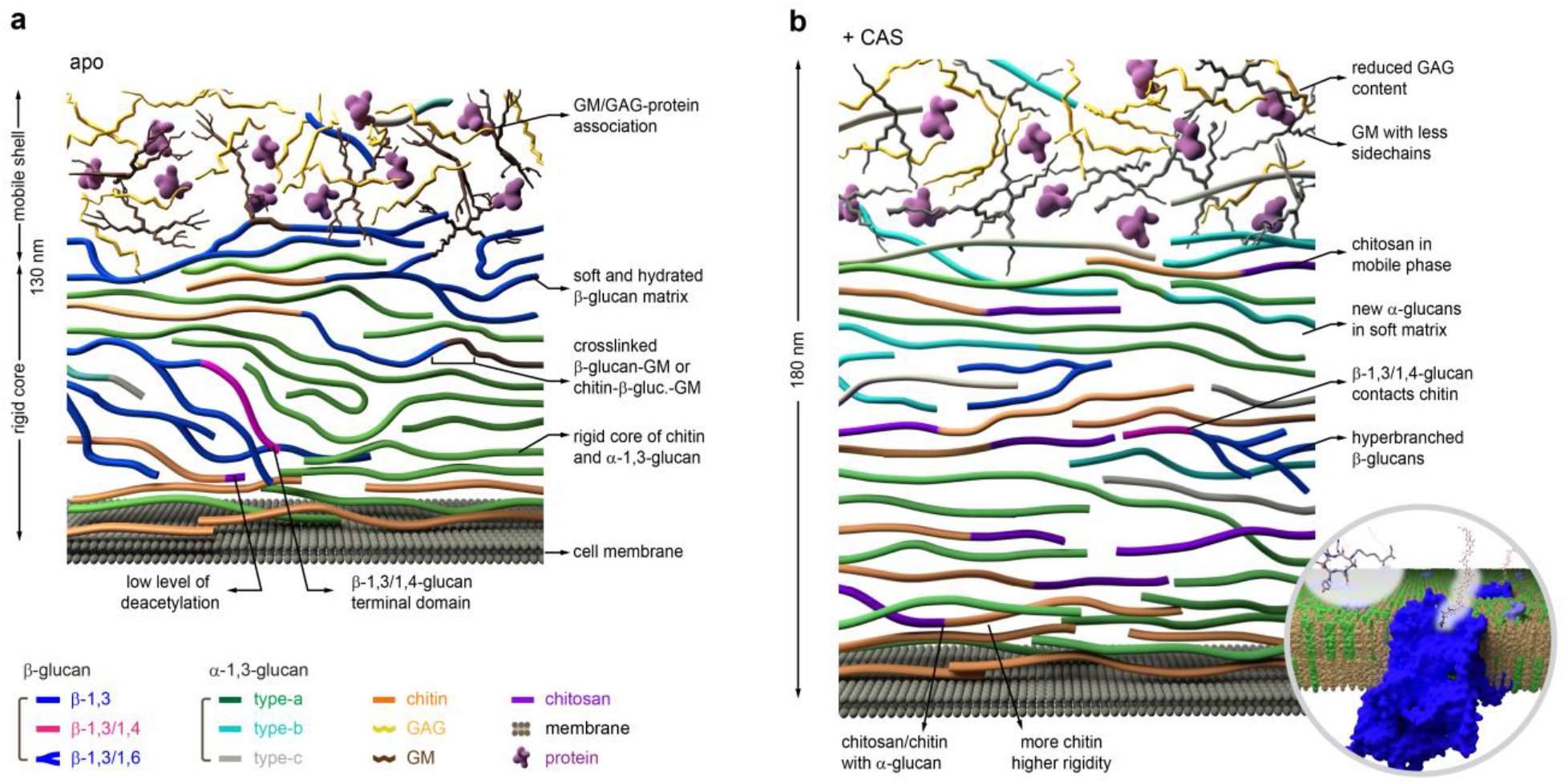
Schematic illustration of cell wall reorganization induced by caspofungin treatment. The illustration integrates NMR observations with the biochemical understanding of *A. fumigatus* cell walls. **a**, Cell walls of untreated 3-day-old *A. fumigatus* mycelial cell walls formed by various biopolymers outside the cell membrane. Molecules shown in the figure include chitin (orange), α-1,3-glucan (green for type-a, cyan for type-b, and grey for type-c), β-1,3-glucan (blue, linear), β-1,3/1,6-glucan (blue, branched), β-1,3/1,4-glucan (magenta), chitosan (purple strands), GAG (yellow), GM (brown), cell membranes (dark grey), and proteins (purple particles). **b**, Following caspofungin treatment, the amount of β-1,3-glucan is substantially reduced while the content of chitin, chitosan, and the two minor forms of α-1,3-glucans increased. The physical contacts among the remaining biopolymers also increased. A zoom-in region of the lipid bilayers of the cell membrane shows β-1,3-glucan synthase. Illustrative models are based on data on cell wall thickness, molecular composition, and intermolecular contacts. The plots may not be strictly to scale.

This high-resolution ssNMR analysis has delineated eight structural mechanisms underlying the multifaceted reconfiguration of the cell wall architecture triggered by caspofungin exposure (Fig. 5b). Firstly, the soft β-1,3-glucan matrix vanishes, resulting in the subsequent removal of the covalently linked polysaccharide cores, such as chitin-β-1,3-glucan-GM and β-1,3-glucan-GM, which are commonly found in untreated cells^19^. Secondly, the remaining β-glucan was restructured, with an increased rate of β-1,6-branching. The β-1,3/1,4-glucan linear domains became physically associated with chitin. Thirdly, chitin biosynthesis is enhanced to reinforce the rigidity of the cell wall while maintaining the polymorphic nature of the chitin structure and its association with other forms. Fourthly, approximately one-third of chitin molecules undergo deacetylation, leading to the formation of partially disordered chitosan that are well-integrated into the physically compacted chitin-α-glucan complex. Chitosan can potentially exist either as a component of partially deacetylated chitin or as individual fully deacetylated chain, or even both. Fifthly, two additional forms of semi-dynamic α-1,3-glucans emerge, namely the type-b and type-c allomorphs, exhibiting a wide distribution across both rigid and mobile phases with altered helical screw conformations. These two forms of α-1,3-glucans constitute one-third of the mobile phase with the cell wall, thereby counterbalancing the lack of β-1,3-glucans to re-establish the soft matrix. Sixthly, the cell wall surface now contains almost equal proportions of GM and GAG. The reduction in exopolysaccharide GAG content, along with its constituent monosaccharide GalN, which possesses cationic properties at the fungal cell’s pH, inevitably weakens the adhesive characteristics of the fungal cell wall. GM also exhibits reduced content of Gal*f* sidechains in mycelia treated with the drug. Seventhly, the cell wall exhibits compromised water retention due to the loss of β-glucan, which is key to maintaining water association within the cell wall. Lastly, the nanoscale thickness of the cell walls increases from 130 nm to 180 nm for better protection.

The β-1,3-glucan synthase is a membrane-embedded enzyme complex comprising at least two subunits, with fks1 being the catalytically active domain^54,55^. Caspofungin acts as a non-competitive inhibitor of the Fks1 subunit, leading to reduced production of β-1,3-glucan. However, *A. fumigatus* retains a functional cell wall structure following treatment with β-glucan inhibitors. The stability of drug-treated cell walls is upheld by numerous physical interactions involving different chitin allomorphs and the interactions between chitin and α-1,3-glucans, which constrain molecular movements and restrict water accessibility within the cell wall. This observation also explains the heightened caspofungin susceptibility of *Candida* species, which possess a skeletal framework consisting solely of β-glucan and chitin while lacking α-1,3-glucan in the rigid phase of the cell wall (**Extended Data Fig. 6**)^33,56^. Eliminating β-1,3-glucan from *Candida* species could potentially lead to a severely impaired cell wall structure, where chitin became the only component responsible for providing mechanical support^56,57^. Thus, structural complexity plays a crucial role in facilitating coordinated changes among cell wall components for antifungal resistance.

As β-glucan has been recognized as a key polysaccharide involved in covalently linking various components^58^, the success of the cell walls devoid of β-1,3-glucan holds two significant implications. First, it suggests that other polysaccharides can substitute for the structural role of β-1,3-glucan, enabling the survival of those fungi with highly intricate cell walls. Second, it emphasizes the importance of packing interactions and physical properties, such as stiffness and water contact, in stabilizing the biopolymer assembly and supporting the cell wall’s function.

Genomic and imaging studies on the clinically important caspofungin paradoxical effect (CPE), characterized by reduced antifungal activity and increased fungal growth rate at high drug concentrations, have proposed potential changes in cell wall structure linked to upregulated chitin synthesis and restored β-glucan synthesis^37,38,59^. Our findings have revealed that remarkable structural dynamics at both the chemical and nanoscale levels are crucial for fungi to withstand stress, including exposure to caspofungin.

These discoveries have advanced our comprehension of cell wall structural integrity and have identified promising avenues for the improvement of antifungal therapies using echinocandins. One approach involves refining existing antifungal compounds to specifically target the modified structures of residual β-glucans and/or the newly formed dynamic α-glucan counterparts that are being used by the fungus as substitutes for the missing β-glucans. Additionally, combination of echinocandins with drugs blocking chitosan or α-1,3-glucan biosynthesis will minimize the compensatory effects observed among fungal pathogens and promote antifungal efficiency.

## Methods

### Preparation of ^13^C, ^15^N-fungal materials

*A. fumigatus* (strain RL 578) was grown in Yeast Extract Peptone Dextrose (YPD) for seven days at 30 °C. Liquid cultures were prepared using a modified minimum medium containing 10.0 g/L of ^13^C-glucose and 6.0 g/L of ^15^N-sodium nitrate for uniformly labeling the fungal material^60^. The medium was adjusted to pH 6.5. Four batches of *A. fumigatus* cultures were prepared in parallel using two different time durations (3 d and 10 d) with and without caspofungin (2.5 µg/mL; above the minimum inhibitory concentration)^61^. These cultures were cultivated in 100 mL liquid medium using 250 mL Erlenmeyer flasks in a shaking incubator (210 rpm) at 30 °C. The fungal pellets were collected by centrifugation at 7000 g for 20 min. The harvested pellets were then washed thoroughly using 10 mM phosphate buffer (PBS, pH 7.4) to remove small molecules and reduce ion concentration. For each sample, approximately 100 mg of the whole-cell material was packed into a 4-mm magic-angle spinning (MAS) rotor for ssNMR characterization. Another 30 mg of material was packed into a 3.2-mm sapphire rotor for DNP experiments. The native hydration level of the fungal cell was fully retained. In addition, *A. sydowii* was grown in minimum media with ^15^N-ammonium sulfate for 7 days at 28 °C, and *C. albicans* was grown in yeast nitrogen-based (YNB) without amino acids and ammonium sulfate with ^15^N-ammonium sulfate for 3 days at 30 °C. ^13^C-glucose was used for both cultures as the carbon source.

### TEM imaging

The harvested *A. fumigatus* mycelial cultures were fixed with 2.5% (vol/vol) glutaraldehyde and 2% (wt/vol) paraformaldehyde in 0.1 M PBS buffer (pH 7.4). The suspensions were centrifuged and embedded in 3% agarose gel. Subsequently, the samples were rinsed with 0.1 M PBS (pH 7.4), 0.05 M glycine, and then post-fixed with 2% osmium tetroxide (OsO_4_). The samples were rinsed thrice using deionized water. Acetone (50%, 70%, 80%, 90%, and 100 %) and propylene oxide were used to dehydrate the samples in two 15 minutes cycles. Finally, a series of propylene oxide: Epon was used for infiltration. Ultrathin sections of the resulting samples were taken for TEM assessments. To increase the contrast during imaging, 1% uranyl acetate *En Bloc* staining was used. All images were taken on perpendicular cross-sections of hyphae using a JEOL JEM-1400 electron microscope.

### Cell wall fractionation and carbohydrate analysis

For carbohydrate analysis, mycelia were disrupted with 1 mm glass beads in 0.2 M Tris-HCl buffer, pH 8.0 with a fast-prep cell disruptor (MP-Bio). Crude cell wall fraction was collected by centrifugation (4,500 × g, 10 min), washed three times with distilled water, and then fractionated into alkali-insoluble and alkali-soluble as previously described^20,62^. Hexose and hexosamine were quantified by colorimetric and chromatographic assays^20,62^. Neutral monosaccharides were analyzed by gas-liquid chromatography (GC) as alditol acetates obtained after hydrolysis (4 N trifluoroacetic acid, 100 °C, 4 h). Derivatized monosaccharides were separated and quantified on a DB5 capillary column (25 m x 0.32 mm, SGE) using a Perichrom GC apparatus (carrier gas, 0.7 bar helium; temperature program, 120-180 °C at 2 °C/min and 180-240 °C at 4 °C/min). To quantify the content of hexosamine, cell wall fractions were hydrolyzed with 6 N HCl at 100°C for 6 h and were analyzed by high performance anion exchange chromatography (HPAEC) with a pulsed electrochemical detector and an anion exchange column (CarboPAC PA-1, 4.6 x 250 mm, Dionex) using 18 mM NaOH as mobile phase at a flow rate of 1 mL/min; glucosamine and galactosamine were used as standards.

To quantify β-1,3-glucan and α-1,3-glucan in the cell wall, the alkali-insoluble fraction and the water-soluble supernatant fraction were submitted to enzymatic digestions. Fractions (1 mg/mL) were incubated with recombinant β-1,3-glucanase (LamA from *Thermotoga neapolitana*) and the mutanase from *Trichoderma harzianum*, respectively, in 20 mM sodium acetate buffer (pH 5.5) at 37 °C for 24 h. The reducing sugar released after enzyme digestion were quantified by the 4-hydroxybenzhydrazide (PABA) assay. To quantify β-1,6-branching in β-1,3-glucan and β-1,3/1,4-glucan, the enzyme digests of different cell wall fractions were subjected to HPAEC using a CarboPAC PA-1 column (4.6 x 250 mm, Dionex) at a flow rate of 1 mL/min; eluent A was 50 mM NaOH, and eluent B was 0.5 M NaOAc in 50 mM NaOH. The elution gradient was: 0-2 min, isocratic 98% A:2% B; 2-15 min, linear gradient from 98% A:2% B to 65% A:35% B; 15-35 min, linear gradient from 65% A:35% B to 30% A:70% B, followed by 100% B for 3 min. Glycosidic linkages were investigated by methylation of cell wall fractions followed by GC-MS analysis as previously described^62,63^. The amount of cell wall galactomannan was estimated from the mannose and galactose content of the alkali-insoluble fraction.

### SsNMR experiments for structural analysis

NMR experiments were conducted on a Bruker Avance 400 MHz (9.4 Tesla) spectrometer and an 800 MHz (18.8 Tesla) using 4-mm and 3.2-mm MAS HCN probes, respectively. All experimental data, except those for MAS-DNP, were collected under 10-13.5 kHz MAS at 290 K. ^13^C chemical shifts were externally referenced to the adamantane CH_2_ peak at 38.48 ppm on the tetramethylsilane (TMS) scale. ^15^N chemical shifts were referenced to the liquid ammonia scale either externally through the methionine amide resonance (127.88 ppm) of the model tri-peptide N-formyl-Met-Leu-Phe-OH or using the ratio of the gyromagnetic ratios of ^15^N and ^13^C^64^. Typical radiofrequency field strengths, unless specifically mentioned, were 80-100 kHz for ^1^H decoupling, 62.5 kHz for ^1^H hard pulses, 50-62.5 kHz for ^13^C, and 41 kHz for ^15^N. The experimental conditions were listed in **Supplementary Table 9**.

One-dimensional (1D) ^13^C spectra were obtained using different polarization methods to selectively detect the rigid and mobile components of the fungal molecules. The rigid components were detected by 1D ^13^C cross-polarization (CP) using 1-ms contact time. Quantitative detection and mobile components located inside the cell wall were measured by 1D ^13^C direct polarization (DP) using short (2-s) and long (35-s) recycle delays, respectively. 2D ^13^C-^13^C 53-ms CORD homonuclear correlation spectra were obtained to detect intramolecular cross-peaks and 2D ^15^N-^13^C N(CA)CX heteronuclear correlation spectra were obtained to detect amide signals of chitin^65^. To measure N(CA)CX, 0.6-ms ^1^H-^15^N CP, 5-ms ^15^N-^13^C CP contact times and 100-ms DARR mixing time were used. The 2D DP refocused J-INADEQUATE spectra were obtained to detect mobile components with through bond connectivity. The assigned ^13^C and ^15^N signals were documented in **Supplementary Table 10**.

To determine the water accessibility of polysaccharides, water-edited 2D ^13^C-^13^C correlation spectra were obtained^66,67^. The experiment was initiated with ^1^H excitation followed by a ^1^H-T_2_ filter of 1.2-ms × 2, which eliminated 97% of the polysaccharide signals but retained 80% of water magnetization. The water magnetization was then transferred to the polysaccharide using a 4-ms ^1^H mixing period and then transferred to ^13^C through a 1-ms ^1^H-^13^C CP for site-specific detection. A 50-ms DARR mixing period was used for both the water-edited spectrum and a control 2D spectrum showing the full intensity. The relative intensity ratio between the water-edited spectrum and the control spectrum was quantified for all cell walls to reflect the relative extent of hydration (**Supplementary Table 7**). The intensities were pre-normalized by the number of scans of each spectrum. Hydration maps were generated using a python script integrated with nmrglue and matplotlib packages^68,69^.

Polysaccharide dynamics were probed using ^13^C spin-lattice (T_1_) relaxation, which was examined using a series of 2D ^13^C-^13^C correlation spectra with different z-filter durations (0-s, 0.1-s, 1-s, 3-s, and 9-s)^70^. The absolute intensity of each peak was quantified and then normalized by the number of scans. Relaxation data were fit using a single exponential function to obtain ^13^C-T_1_ time constants (**Supplementary Table 8**).

### MAS-DNP sample preparation and experiments

The stock solution containing 10 mM AMUPol was freshly prepared using a d_8_-glycerol/D_2_O/H_2_O (60/30/10 Vol%) solvent mixture, typically named DNP-matrix^71^. Around 30 mg of ^13^C,^15^N-labeled mycelial cells were mixed with 50 µL of stock solution and gently ground using a mortar and pestle for 10-15 min, which allows radicals to penetrate the porous cell walls. The material was then transferred to a 3.2-mm sapphire rotor for the DNP measurement. All DNP experiments were performed on a 600 MHz/395 GHz 89 mm-bore MAS-DNP spectrometer equipped with a gyrotron microwave source^72^. All spectra were measured using a 3.2-mm HCN probe under 8 kHz MAS frequency at 92 K. The microwave power was set to 12 W at the probe base. The enhancement factor of NMR sensitivity with and without microwave irradiation (ε_on/off_) was 17–32 for these samples. The buildup time varied between 5.2–13-s.

2D ^13^C-^13^C 100-ms PDSD were measured to detect intra-molecular correlations, followed by 20-ms PAR measurement to detect both intra- and inter-molecular correlations in carbohydrates of each cell wall^73^. The ^13^C and ^1^H irradiation frequencies were 56 kHz and 53 kHz during the PAR mixing. 2D ^15^N-^13^C N(CA)CX spectra were measured using the double-CP sequence^74^, with 1.0 ms contact time for the ^1^H-^15^N CP and 4.0 ms for the ^15^N-^13^C CP. In these ^15^N-^13^C experiments, ^13^C-^13^C mixing was achieved using a PDSD period of either 0.1 s for detecting intramolecular cross peaks or 3.0 s for detecting both intra- and inter-molecular cross peaks. The resonance assignment under DNP condition was presented in **Extended Data Fig. 4a-d**., with the chemical shifts documented in **Supplementary Table 11**. The identified long-range intermolecular cross peaks were listed in **Supplementary Table 12**.

### Construction of atomic models for MD simulation

We first assembled model α- and β-glucans using CHARMM-GUI Glycan Modeler^75,76^. The degree of polymerization (D.P.) of each polysaccharide was set to six monomers, which allowed us to keep the total system size of the whole system under 200,000 atoms as detailed later. Although the D.P. of the polysaccharide in the native fungal cell wall is significantly higher, the current atomistic model allows us to evaluate the structure-based polymer interactions. The CHARMM-GUI Glycan Modeler then assembled the short linear polysaccharides, α-1,3-glucan, β-1,3-glucan, and β-1,3/1,4-glucan as well as the branched β-1,3/1,6-glucan (**Supplementary Fig. 7a**). Similarly, short chitin polymers were generated using the plugin chitin_builder^77^, within visual molecular dynamics (VMD 1.9.4a55)^78^. Chitin and chitosan are copolymers of 2-acetamido-2-deoxy-D-glucose (N-acetyl glucosamine, GlcNAc), and 2-amino-2-deoxy-D-glucose (glucosamine, GlcNH_2_), connected by β-1,4 glycoside linkages^79^. Chitin and chitosan might exist as a copolymer, with a variable degree of N-acetylation. In addition to the copolymer in our molecular system, we created separate polymers for chitin and chitosan to accurately track the individual interactions specific to each of the two polysaccharides. Since glucosamine in the existing CHARMM carbohydrate force field carries a charge^80,81^, we introduced a patch to modify the topology and convert glucosamine into a monosaccharide unit of chitosan by deleting hydrogen and reparametrizing the charges. The final system has two copies of each polysaccharide, which were solvated in a box of water using the solvate plugin in VMD 1.9.4a55, with 5 mM of NaCl added to the hydrated neutralized model using the autoionize plugin to VMD 1.9.4a55^78^. The atomistic model consists of 120,972 atoms after adding solvent and ions (**Supplementary Fig. 7b**).

### MD simulation and analysis of molecular trajectories

All-atom classical MD simulations for the models illustrated in **Figure 4e, f** and **Supplementary Figure 7** were initially minimized using NAMD 2.14, to prepare for production simulations using the GPU resident integrator in NAMD 3.0a9. The simulation employed the CHARMM36 force fields for computing the interactions between carbohydrates^80,81^, water^82^, and ions^83^. Following CHARMM36 force field, we used a 12 Å cutoff and a force-switching function after 10 Å. Long-range electrostatics are treated using the particle mesh Ewald with a 1.2 Å grid spacing^84^. To enable a 2 fs timestep between force evaluations, covalent bonds involving hydrogen atoms were treated using the RATTLE algorithm^85^. The Langevin thermostat was set to maintain a temperature of 300 K using a 5 ps^−1^ damping coefficient^86^. A microsecond-long MD simulation was performed for an NPT ensemble, using a Langevin isotropic barostat that was set to maintain 1 atm pressure^87^.

The simulation trajectory was analyzed using VMD 1.9.4a55, which was enabled with Python 3.10.12 and Numpy 1.24.2^88^. In-house scripts were prepared in Tcl 8.6 and Python 3.10.12 programming software to calculate and quantify the intermolecular interactions between polysaccharides using the coordination number collective variable^89^. The coordination number (C_ij_) measures the pairwise contact between two atoms within a given polymer chain. This was implemented using a switching function defined as 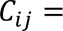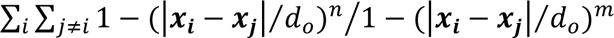. Short- and long-range interactions were defined using the cutoff parameter, which used the switching distance to define an interatomic contact, for all d << d_0_. Exponents n and m are set to 10 and 120, to steepen the switching profile. The equation for the coordination number is illustrated in Supplementary Figure 7 for short-range and long-range cutoff 5 and 7 Å with the exponents n =10 and m = 120. A contact map illustrating the polysaccharide interactions within 5 Å from each other was computed to probe the short-range interactions in the fungal cell wall (Fig. 4g). All system preparation, molecular simulation input and analysis script are made publicly available on Zenodo.

### Reporting summary

Further information on research design is available in the Nature Research Reporting Summary linked to this article.

### Data Availability

All NMR spectra and biochemical data that support the findings of this study are provided in the article, Extended Data, Supplementary Information, and Source Data files. The topspin NMR datasets of 64 ssNMR spectra collected on fungal cell walls are available in the public repository Zenodo under the DOI number: https://doi.org/10.5281/zenodo.8226836. The source data underlying Figures 1c, 1d, 1g, 3b, 3c, and Supplementary Figure 4b, 5c, and 5d are provided as a Source Data file.

### Code Availability

All MD input data, which includes topology, force field parameters and input scripts that support the findings of this research will be made publicly available via Zenodo under the DOI number https://doi.org/10.5281/zenodo.8226836.

### Supplementary Information

Extended Data Fig. 1-6, Supplementary Movie 1-5, Supplementary Figs. 1-7, Supplementary Tables 1-12.

## Supporting information

Movie S1. Fungal cell wall interactions

Movie S2. Alphaglucan chitin chitosan interactions

Movie S3. Betaglucan chitin chitosan interactions

Movie S4. Chitin chitosan interactions

Movie S5. Chitin stacking

## Acknowledgments

This work was supported by the National Institute of Health (NIH) grant AI173270. The National High Magnetic Field Laboratory is supported by the National Science Foundation through NSF/DMR-1644779 and 2128556, and the State of Florida. The MAS-DNP system at NHMFL is funded in part by NIH S10 OD018519, P41 GM122698, and RM1 GM148766. The modeling work was supported in part through computational resources and services provided by the Institute for Cyber-Enabled Research at Michigan State University. D.S. and J.V.V acknowledge support from DE-FG02-91ER20021 from the U.S. Department of Energy, Office of Basic Energy Sciences. The authors thank Drs. Ivan Hung and Zhehong Gan for technical assistance.

## Authors Contribution

M.C.D.W. prepared fungal samples and conducted the NMR experiments. F.M.V. conducted MAS-DNP experiments. M.C.D.W and I.G. analyzed the NMR, DNP, and TEM data. J.V.V. and D.S. performed MD simulations. T.F. and J.P.L. performed the chemical analysis of fungal cell walls. All authors contributed to the writing of the manuscript. P.W. and T.W. designed and supervised the project.

## Extended Data Figures

**Extended Data Fig. 1.**
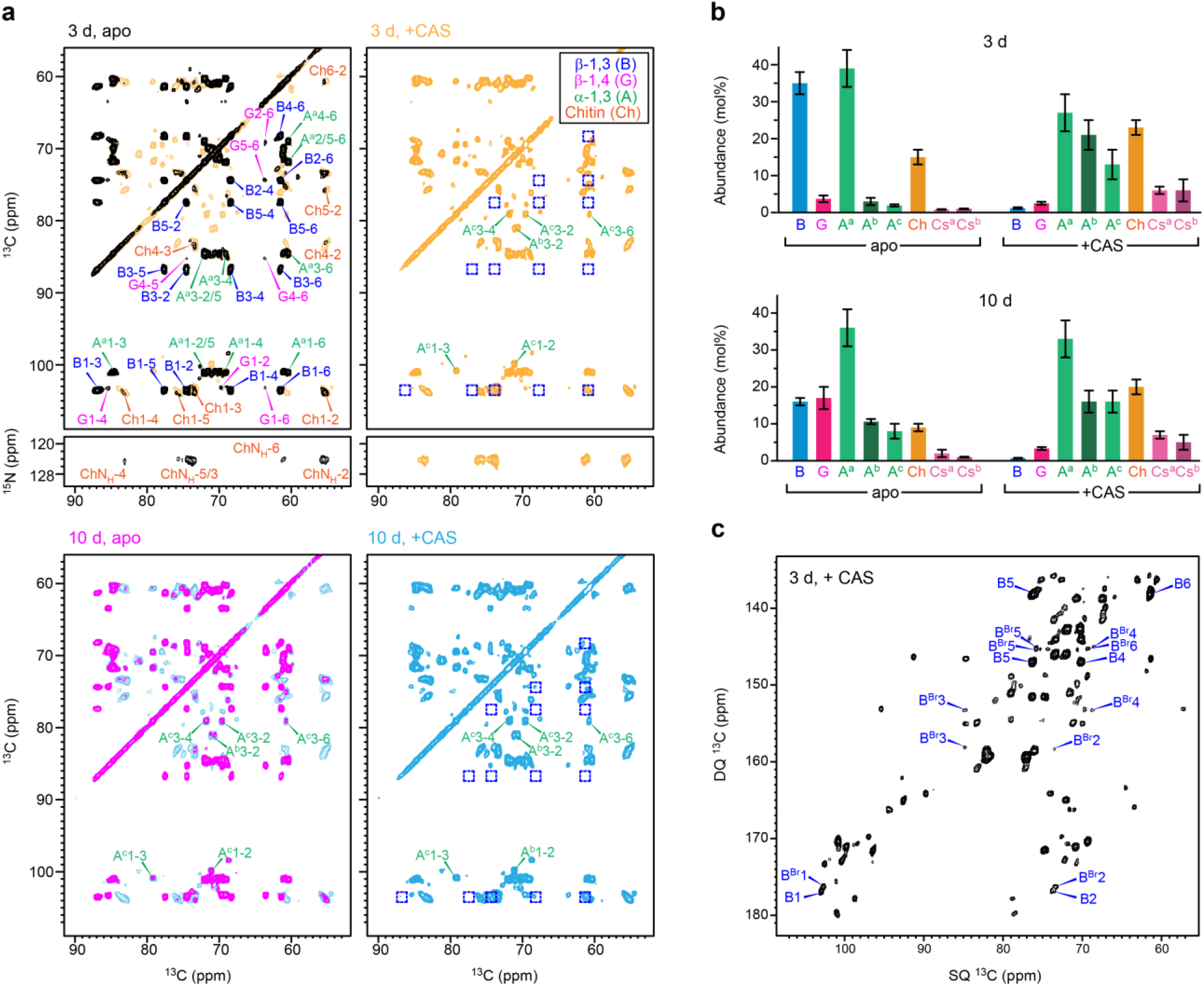
Effect of caspofungin and culture duration on cell wall composition. **a**, 2D ^13^C-^13^C/^15^N spectra of 3-day-old (top) and 10-day-old (bottom) *A. fumigatus* cell walls without and with caspofungin (CAS). The spectra were measured using CP and 53-ms CORD mixing for detecting chitin and glucans in the rigid portion of *A. fumigatus* cell walls. The ^15^N-^13^C correlation spectra were measured using NCACX sequence, showing correlations between chitin N_H_ and carbons. In the first column, the spectra of drug-treated and apo samples were overlaid for comparison. Abbreviations are used for resonance assignment and different polysaccharide signals are color-coded. The missing peaks of β-1,3-glucan linkage in drug-treated cell walls is marked using blue squares. **b**, Molar composition of cell wall polysaccharides. The compositional data was obtained by analysis of the intensities of resolved cross peaks in 2D spectra. Error bars represent standard error derived from the intensities of the different cross-peaks for each type of polysaccharides. **c**, 2D ^13^C DP J-INADEUQATE spectrum of 3-day-old drug-treated *A. fumigatus* resolves the signals from the linear chain (B) and the branching points (B^Br^) of β-glucans.

**Extended Data Fig. 2.**
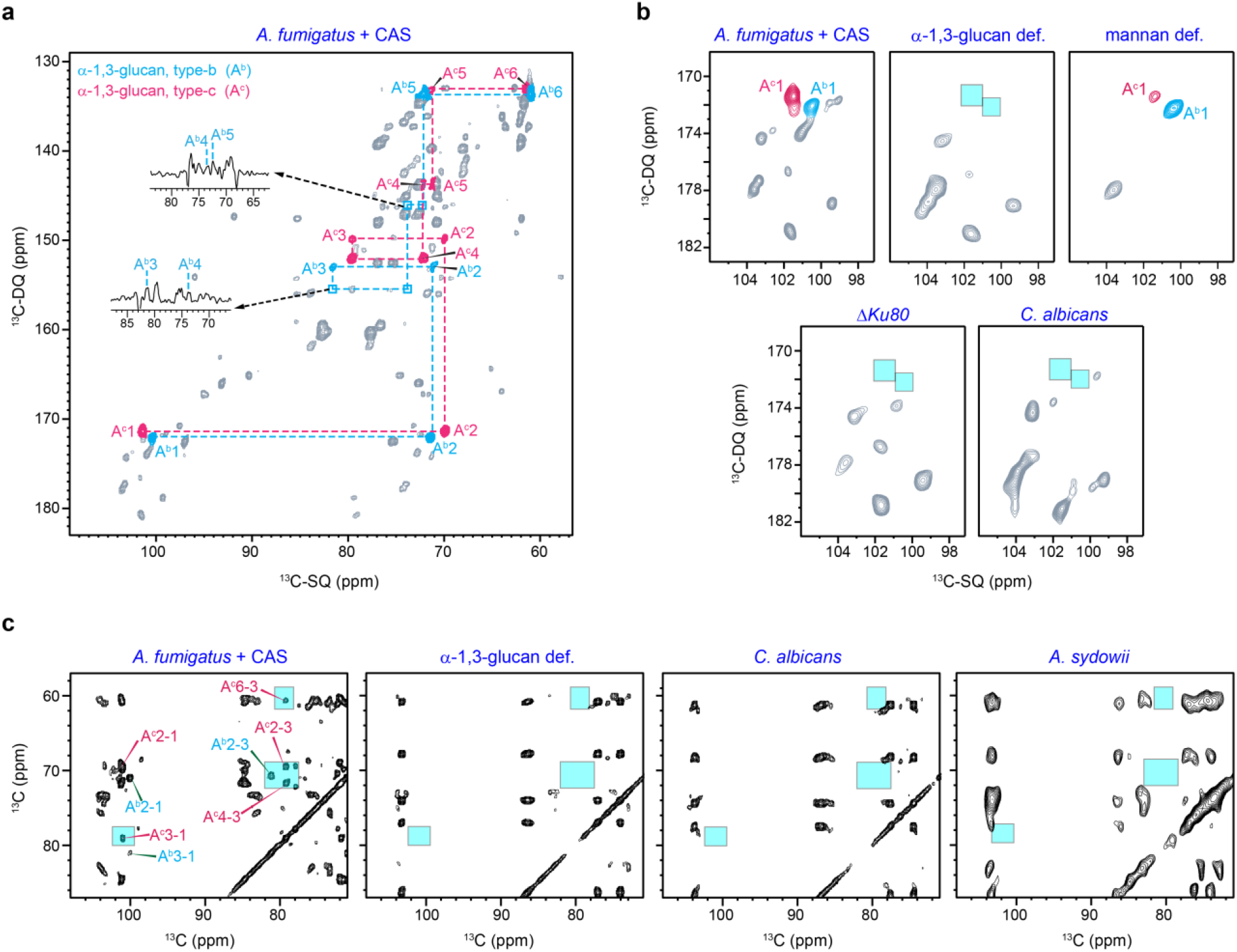
Identification of two minor forms of α-1,3-glucans. **a**, Carbon connectivity of type-b (A^b^; cyan) and type-c (A^c^; magenta) forms of α-1,3-glucans in the caspofungin-treated *A. fumigatus* identified in 2D ^13^C DP J-INADEUQATE spectra. Cross-sections are shown for signals that are weak and below the plotted contour level. **b**, Comparison of the carbon-1 regions of DP J-INADEUQATE spectra confirms the identity of the two minor forms of α-1,3-glucans and their presence in the mobile portion of cell wall polysaccharides. The type-b and type-c forms of α-1,3-glucans are resolved in the drug-treated cell walls of wild-type *A. fumigatus* (RL578) but absent in the α-1,3-glucan-deficient mutant, thus confirming the identity and assignment of these signals. Cyan boxes are used to highlight the missing signals of the minor forms of α-1,3-glucans. The signals are also absent in the mannan-deficient mutant, thus excluding the attribution from the manna polymers that dominate this type of spectra. The signals are absent in a widely used model strain Δ*akuB^KU^*^80^ of *A. fumigatus*, and in *Candida albicans*, evidencing their unique abundance in the caspofungin-treated A. fumigatus. **c**, Confirmation of the presence of type-b and type-c forms of α-1,3-glucans in the rigid portion of caspofungin-treated A. fumigatus cell walls as detected using 53-ms CORD spectrum. The corresponding signals are confirmed to be absent in α-1,3-glucan-deficient mutant of A. fumigatus, as well as in *C. albicans*, and *A. sydowii.* All 2D ^13^C-^13^C correlation spectra were collected on an 800 MHz ssNMR spectrometer, except the spectrum of *A. sydowii*, which was measured on a 400 MHz NMR instrument.

**Extended Data Fig. 3.**
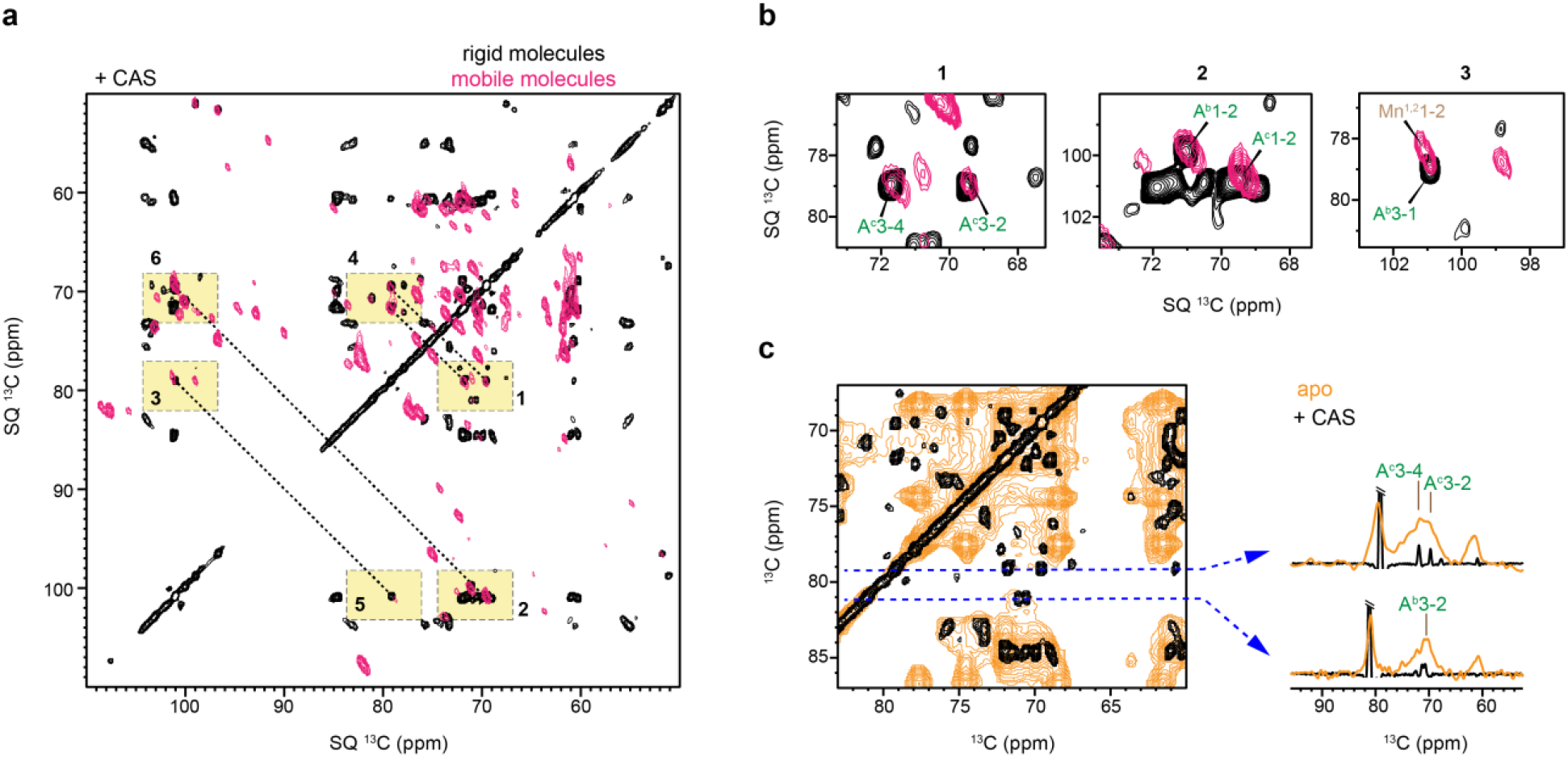
Distribution of minor forms of α-1,3-glucans in dynamically distinct domains. **a**, Overlay of two 2D ^13^C-^13^C correlation spectra of 3-day-old *A. fumigatus* treated with caspofungin: the CP-based 53-ms CORD spectrum (black) selecting rigid molecules and the sheared DP-based refocused J-INADEQUATE spectrum (magenta) selecting mobile components. Both spectra exhibit signals of the type-b (A^b^) and type-c (A^c^) of α-1,3-glucans in six highlighted regions. **b**, Zoomed-in regions extracted from panel a, confirming that the type-b and type-c α-1,3-glucan are distributed in both rigid and mobile fractions of drug-treated *A. fumigatus* cell walls. **c**, Type-c of α-1,3-glucan is also present in the drug-free *A. fumigatus* cell walls but with low abundance, which is shown by the overlay of 53-ms CORD spectra of 3-day-old cell wall with (orange) and without (black) caspofungin. The cross-sections extracted at 81 ppm and 79 ppm show the signals of type-b and type-c forms, respectively. The spectrum of apo sample is processed with more line-broadening to show the weaker signals of these two minor forms of α-1,3-glucans in untreated *A. fumigatus.* All spectra were collected on an 800 MHz NMR under 12 kHz MAS.

**Extended Data Fig. 4.**
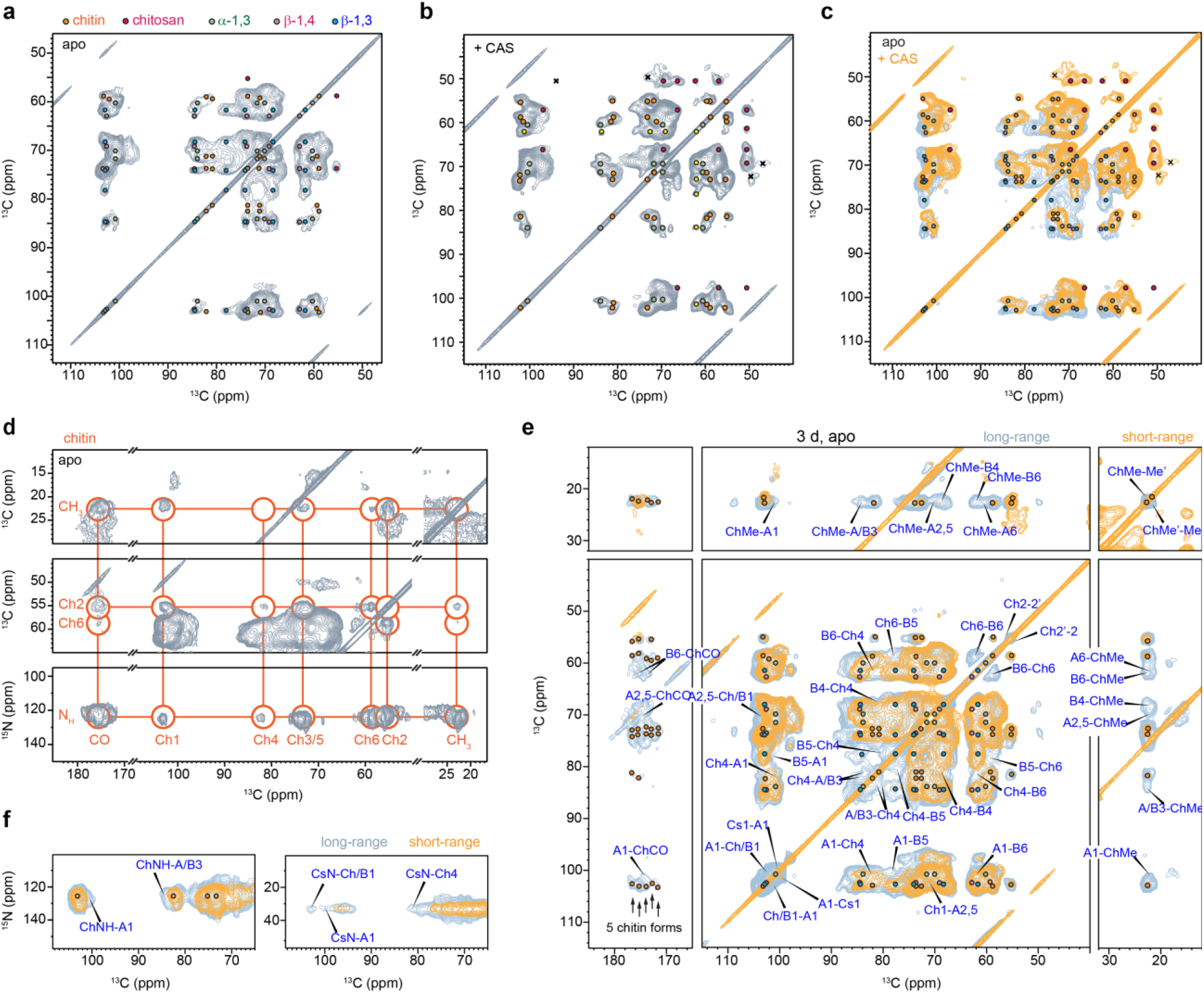
DNP analysis of cell wall polysaccharides and packing interface. Carbohydrate resonance assignment was shown for **a**, apo and **b**, drug-treated 3-day-old samples. The DNP 2D ^13^C-^13^C spectra were measured using 100-ms PDSD mixing. **c**, Overlay of 2D ^13^C-^13^C spectra of 3-day-old apo (grey) and drug-treated (yellow) samples showing the removal of β-1,3-glucan after CAS treatment. **d**, DNP 2D ^15^N/^13^C-^13^C correlation spectra show the carbon and nitrogen connectivity in chitin in the apo sample. **e**, Intermolecular interactions of 3-day-old cell wall detected by overlaying 100-ms PDSD (short range; yellow) and 20-ms PAR (long range; grey) spectra under DNP enhancement. **f,** ^15^N-^13^C correlation spectra of 3-day-old apo sample measured with 100-ms (yellow) and 3-s (grey) ^13^C-^13^C mixing periods. The cross peaks are between chitin -NH-(left panel) and chitosan -NH_2_ (right panel) with carbohydrate carbon sites. All spectra were collected on a 600 MHz/395 GHz MAS-DNP spectrometer.

**Extended Data Fig. 5.**
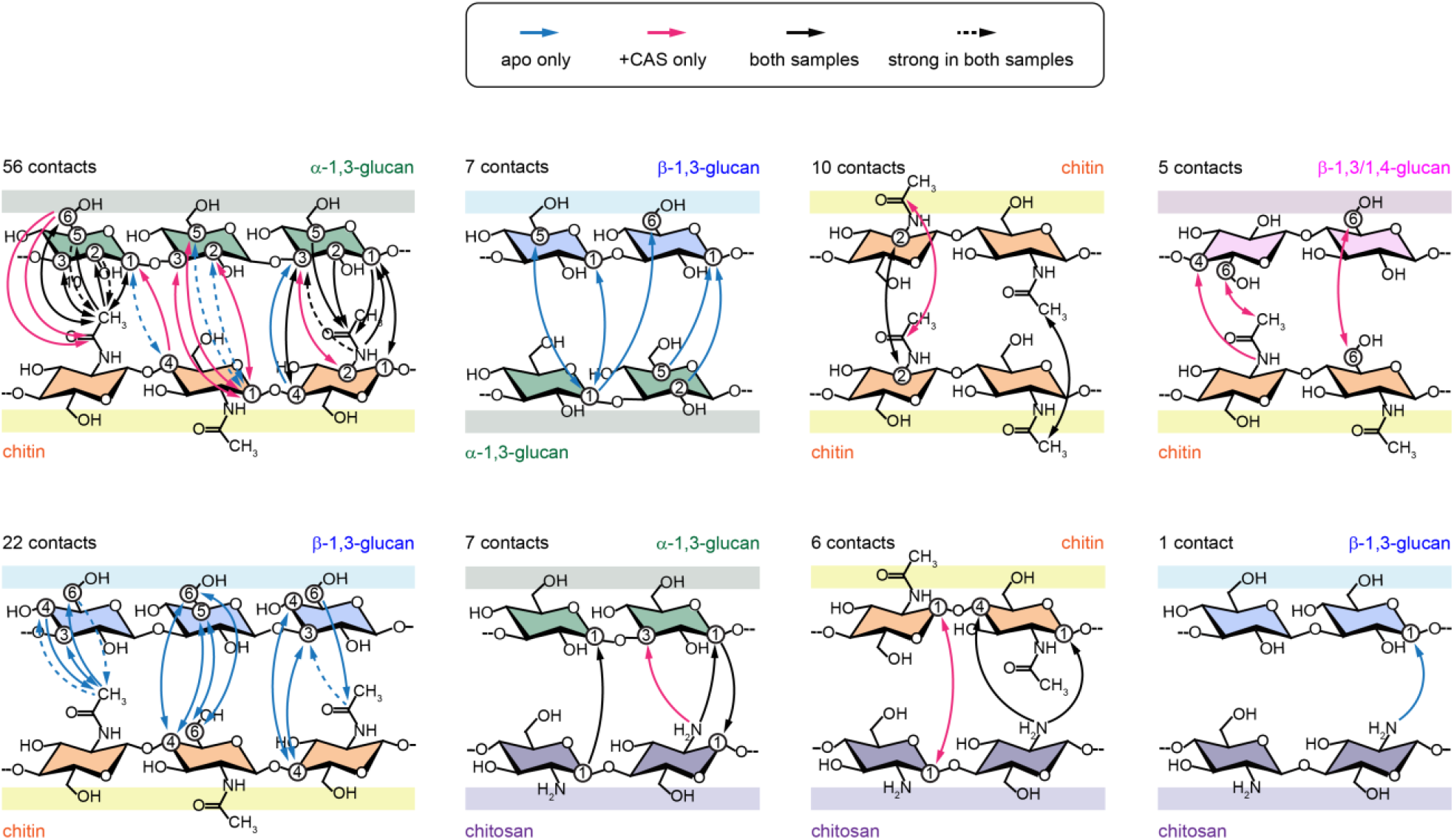
Structural summary of NMR-observed intermolecular contacts. The 114 intermolecular cross peaks identified in 3-day-old *A. fumigatus* cell walls are categorized based on polysaccharide species at the packing interface. Interactions unique to the apo sample are depicted by blue solid lines, those exclusive to drug-treated samples by magenta solid lines, and shared interactions by black solid lines. Strong cross peaks observed in short-mixing 2D correlation spectra and present in both apo and drug-treated samples are indicated by black dashed lines. Arrowheads denote the direction of polarization transfer between carbon and nitrogen sites, encompassing both one-way (one cross peak) and bidirectional transfers (two cross peaks). The number of cross peaks observed between each pair of polysaccharides is indicated in each figure panel.

**Extended Data Fig. 6.**
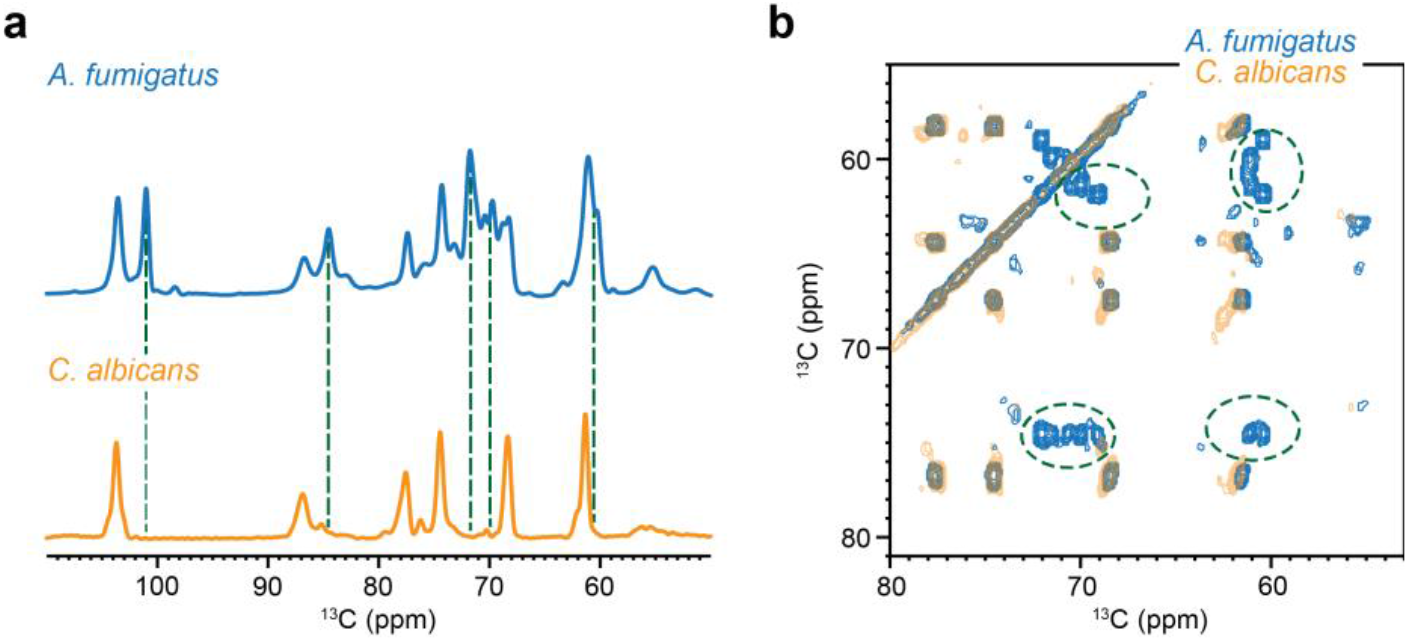
Simpler Rigid Domain in *C. albicans* Cell Wall Compared to *A. fumigatus*. **a**, 1D ^13^C CP spectrum of *C. albicans* (yellow) exhibiting fewer peaks than *A. fumigatus* (blue). Green dash lines indicate the positions of α-1,3-glucan peaks absent in *C. albicans*. **b**, Overlay of two 2D ^13^C-^13^C 53 ms CORD spectra collected on *A. fumigatus* (blue) and *C. albicans* (yellow). Signals of α-1,3-glucans are highlighted in green dash line circles. All spectra were measured on 800 MHz NMR spectrometer at 12-13.5 kHz MAS.

## References

1 Brown, G. D. et al. Hidden killers: human fungal infections. Sci Transl Med 4, 165rv113 (2012).

2 Alanio, A. et al. Azole resistance of Aspergillus fumigatus in immunocompromised patients with invasive aspergillosis. Emerging Infect. Dis. 22, 157 (2016).

3 Perlin, D. S. Current perspectives on echinocandin class drugs. Future Microbiol 6, 441–457 (2011).

4 Chen, S. C. A., Slavin, M. A. & Sorrell, T. C. Echinocandin antifungal drugs in fungal infections. Drugs 71, 11–41 (2011).

5 Garcia-Vidal, C., Viasus, D. & Carratala, J. Pathogenesis of invasive fungal infections. Curr. Opin. Infect. Dis. 26, 270–276 (2013).

6 Erwig, L. P. & Gow, N. A. R. Interactions of fungal pathogens with phagocytes. Nat. Rev. Microbiol. 14, 163–176 (2016).

7 Hoenigl, M. Invasive Fungal Disease Complicating Coronavirus Disease 2019: When It Rains, It Spores. Clin. Infect. Dis. 73, e1645–e1648 (2021).

8 Hoenigl, M. et al. COVID-19-associated fungal infections. Nat. Microbiol. 7, 1127–1140 (2022).

9 Ghannoum, M. A. & Rice, L. B. Antifungal agents: mode of action, mechanisms of resistance, and correlation of these mechanisms with bacterial resistance. Clin. Microbiol. Rev. 12, 501–517 (1999).

10 Verweij, P. E., Snelders, E., Kema, G. H. J., Mellado, E. & Melchers, W. J. G. Azole resistance in Aspergillus fumigatus: a side-effect of environmental fungicide use? Lancet Infect Dis 9, 789–795 (2009).

11 Odds, F. C., Brown, A. J. P. & Gow, N. A. R. Antifungal agents: mechanisms of action. Trends Microbiol. 11, 272–279 (2003).

12 Hasim, S. & Coleman, J. J. Targeting the fungal cell wall: current therapies and implications for development of alternative antifungal agents. Future Med Chem 11, 869–883 (2019).

13 SC, D. Stevens DA. Caspofungin. Clin. Infect. Dis. 36, 1445–1457 (2003).

14 Hu, X. et al. Structural and mechanistic insights into fungal β-1,3-glucan synthase FKS1. Nature 616, 190–198 (2023).

15 Reboli, A. C. et al. Anidulafungin versus fluconazole for invasive candidiasis. N. Engl. J. Med. 356, 2472–2482 (2007).

16 Pappas, P. G. et al. Micafungin versus caspofungin for treatment of candidemia and other forms of invasive candidiasis. Clin. Infect. Dis. 45, 883–893 (2007).

17 Latgé, J. P. The pathobiology of Aspergillus fumigatus. Trends Microbiol. 9, 382–389 (2001).

18 Latgé, J. P. Aspergillus fumigatus and aspergillosis. Clin. Microbiol. Rev. 12, 310–350 (1999).

19 Latgé, J. P. The cell wall: a carbohydrate armour for the fungal cell. Mol. Microbiol. 66, 279–290 (2007).

20 Fontaine, T. et al. Molecular organization of the alkali-insoluble fraction of Aspergillus fumigatus cell wall. J. Biol. Chem. 275, 27594–27607 (2000).

21 Bowman, J. C. et al. The antifungal echinocandin caspofungin acetate kills growing cells of Aspergillus fumigatus in vitro. Antimicrob. Agents Chemother. 46, 3001–3012 (2002).

22 Gardiner, R. E., Souteropoulos, P., Park, S. & Perlin, D. S. Characterization of Aspergillus fumigatus mutants with reduced susceptibility to caspofungin. Med. Mycol. 43, S299–S305 (2005).

23 Satish, S. & Perlin, D. S. Echinocandin resistance in Aspergillus fumigatus has broad implications for membrane lipid perturbations that influence drug-target interactions. Microbiol Insights 12, 1178636119897034 (2019).

24 Satish, S. et al. Stress-induced changes in the lipid microenvironment of β-(1, 3)-D-glucan synthase cause clinically important echinocandin resistance in Aspergillus fumigatus. MBio 10, e00779–00719 (2019).

25 Safeer, A. et al. Probing Cell-Surface Interactions in Fungal Cell Walls by High-Resolution 1H-Detected Solid-State NMR Spectroscopy. Chem. Euro. J. 29, e202202616 (2022).

26 Chrissian, C. et al. Solid-state NMR spectroscopy identifies three classes of lipids in Cryptococcus neoformans melanized cell walls and whole fungal cells. J. Biol. Chem. 295, 15083–15096 (2020).

27 Ghassemi, N. et al. Solid-State NMR Investigations of Extracellular Matrixes and Cell Walls of Algae, Bacteria, Fungi, and Plants. Chem. Rev. 122, 10036–10086 (2022).

28 Kang, X. et al. Molecular architecture of fungal cell walls revealed by solid-state NMR. Nat. Commun. 9, 2747 (2018).

29 Latgé, J. P. & Wang, T. Modern Biophysics Redefines Our Understanding of Fungal Cell Wall Structure, Complexity, and Dynamics. mBio 13, e0114522 (2022).

30 Lamon, G. et al. Solid-state NMR molecular snapshots of Aspergillus fumigatus cell wall architecture during a conidial morphotype transition. Proc. Natl. Acad. Sci. USA 120, e2212003120 (2023).

31 Chakraborty, A. et al. A molecular vision of fungal cell wall organization by functional genomics and solid-state NMR. Nat. Commun. 12, 6346 (2021).

32 Hopke, A., Brown, A. J. P., Hall, R. A. & Wheeler, R. T. Dynamic fungal cell wall architecture in stress adaptation and immune evasion. Trends Microbiol. 26, 284–295 (2018).

33 Garcia-Rubio, R., de Oliveira, H. C., Rivera, J. & Trevijano-Contador, N. The fungal cell wall: Candida, Cryptococcus, and Aspergillus species. Front. Microbiol. 10, 2993 (2020).

34 Klis, F. M., Mol, P., Hellingwerf, K. & Brul, S. Dynamics of cell wall structure in Saccharomyces cerevisiae. FEMS Microbiol. Rev. 26, 239–256 (2002).

35 Cowen, L. E. & Steinbach, W. J. Stress, drugs, and evolution: the role of cellular signaling in fungal drug resistance. Eukaryotic Cell 7, 747–764 (2008).

36 Chiou, C. C., Mavrogiorgos, N., Tillem, E., Hector, R. & Walsh, T. J. Synergy, pharmacodynamics, and time-sequenced ultrastructural changes of the interaction between nikkomycin Z and the echinocandin FK463 against Aspergillus fumigatus. Antimicrob. Agents Chemother. 45, 3310–3321 (2001).

37 Ries, L. N. A. et al. The Aspergillus fumigatus CrzA transcription factor activates chitin synthase gene expression during the caspofungin paradoxical effect. MBio 8, e00705–00717 (2017).

38 Loiko, V. & Wagener, J. The paradoxical effect of echinocandins in Aspergillus fumigatus relies on recovery of the β-1, 3-glucan synthase Fks1. Antimicrob. Agents Chemother. 61, e01690–01616 (2017).

39 Fernando, L. D., et al. Structural Organization of the Cell Wall of Halophilic Fungi. Preprint at bioRxiv, DOI: 10.1101/2023.04.15.537024 (2023).

40 Samar, D., Kieler, J. B. & Klutts, J. S. Identification and deletion of Tft1, a predicted glycosyltransferase necessary for cell wall β-1, 3; 1, 4-glucan synthesis in Aspergillus fumigatus. PLoS One 10, e0117336 (2015).

41 Mouyna, I. et al. Glycosylphosphatidylinositol-anchored glucanosyltransferases play an active role in the biosynthesis of the fungal cell wall. J. Biol. Chem. 275, 14882–14889 (2000).

42 Ao, J., Chinnici, J. L., Maddi, A. & Free, S. J. The N-Linked Outer Chain Mannans and the Dfg5p and Dcw1p Endo-α-1,6-Mannanases Are Needed for Incorporation of Candida albicans Glycoproteins into the Cell Wall. Eukaryot. Cell 14, 792–803 (2015).

43 Simmons, T. J. et al. Folding of xylan onto cellulose fibrils in plant cell walls revealed by solid-state NMR. Nat. Commun. 7, 13902 (2016).

44 Kirui, A. et al. Carbohydrate-aromatic interface and molecular architecture of lignocellulose. Nat. Commun. 13, 538 (2022).

45 Temple, H. et al. Golgi-localized putative S-adenosyl methionine transporters required for plant cell wall polysaccharide methylation *Nat*. Plants 8, 656–669 (2022).

46 Ni, Q. Z. et al. High frequency dynamic nuclear polarization. Acc. Chem. Res. 46, 1933–1941 (2013).

47 Mentink-Vigier, F., Akbey, Ü., Oschkinat, H., Vega, S. & Feintuch, A. Theoretical aspects of magic angle spinning-dynamic nuclear polarization. J. Magn. Reson. 258, 102–120 (2015).

48 Koers, E. J. et al. NMR-based structural biology enhanced by dynamic nuclear polarization at high magnetic field. J. Biomol. NMR 60, 157–168 (2014).

49 Fernando, L. D. et al. Structural polymorphism of chitin and chitosan in fungal cell walls from solid-state NMR and principal component analysis. Front. Mol. Biosci., 727053 (2021).

50 Sarkar, D. et al. Diffusion in intact secondary cell wall models of plants at different equilibrium moisture content. Cell Surf. 9, 100105 (2023).

51 Vermaas, J. V. et al. Mechanism of lignin inhibition of enzymatic biomass deconstruction. Biotechnol. Biofuels Bioprod. 8, 217 (2015).

52 N.A.R., G., Latge, J. P. & Munro, C. A. The Fungal Cell Wall: Structure, Biosynthesis, and Function. Microbiol. Spectr. 5, FUNK-0035-2016 (2017).

53 Wessels, J. G. H. Developmental regulation of fungal cell wall formation. Annu. Rev. Phytopathol. 32, 413–437 (1994).

54 Schimoler-O’Rourke, R., Renault, S., Mo, W. & Selitrennikoff, C. P. Neurospora crassa FKS protein binds to the (1, 3) β-glucan synthase substrate, UDP-glucose. Curr. Microbiol. 46, 0408–0412 (2003).

55 Douglas, C. M. et al. The Saccharomyces cerevisiae FKS1 (ETG1) gene encodes an integral membrane protein which is a subunit of 1, 3-beta-D-glucan synthase. Proc. Natl. Acad. Sci. U.S.A. 91, 12907–12911 (1994).

56 Gow, N. A. R. & Lenardon, M. D. Architecture of the dynamic fungal cell wall. Nat. Rev. Microbiol. 21, 248–259 (2022).

57 Stevens, D. A., Espiritu, M. & Parmar, R. Paradoxical effect of caspofungin: reduced activity against Candida albicans at high drug concentrations. Antimicrob. Agents Chemother. 48, 3407–3411 (2004).

58 Aimanianda, V. et al. The Dual Activity Responsible for the Elongation and Branching of β-(1,3)-Glucan in the Fungal Cell Wall. mBio 8, 00619–00617 (2017).

59 Zhao, S. et al. Genomic and Molecular Identification of Genes Contributing to the Caspofungin Paradoxical Effect in Aspergillus fumigatus. Microbiol. Spectr. 10, 00519–00522 (2022).

## References for Methods

60 Kirui, A. et al. Preparation of fungal and plant materials for structural elucidation using dynamic nuclear polarization solid-state NMR. J. Vis. Exp. (2019).

61 McCormack, P. L. & Perry, C. M. Caspofungin. Drugs 65, 2049–2068 (2005).

62 Liu, Z. et al. Conidium Specific Polysaccharides in Aspergillus fumigatus. J. Fungi 9, 155 (2023).

63 Johnson, S. B. & Brown, R. E. Simplified derivatization for determining sphingolipid fatty acyl composition by gas chromatography-mass spectrometry *J*. Chromatogr. 605, 281–286 (1992).

64 Rienstra, C. M. et al. De novo determination of peptide structure with solid-state magic-angle spinning NMR spectroscopy. Proc. Natl. Acad. Sci. U.S.A. 99, 10260–10265 (2002).

65 Baldus, M., Petkova, A. T., Herzfeld, J. & Griffin, R. G. Cross polarization in the tilted frame: assignment and spectral simplification in heteronuclear spin systems. Mol. Phys. 95, 1197–1207 (1998).

66 Ader, C. et al. Structural rearrangements of membrane proteins probed by water-edited solid-state NMR spectroscopy. J. Am. Chem. Soc. 131, 170–176 (2009).

67 White, P. B., Wang, T., Park, Y. B., Cosgrove, D. J. & Hong, M. Water–polysaccharide interactions in the primary cell wall of Arabidopsis thaliana from polarization transfer solid-state NMR. J. Am. Chem. Soc. 136, 10399–10409 (2014).

68 Helmus, J. J. & Jaroniec, C. P. Nmrglue: an open source Python package for the analysis of multidimensional NMR data. J. Biomol. NMR 55, 355–367 (2013).

69 Hunter, J. D. Matplotlib: A 2D graphics environment. Comput. Sci. Eng. 9, 90–95 (2007).

70 Wang, T., Williams, J. K., Schmidt-Rohr, K. & Hong, M. Relaxation-compensated difference spin diffusion NMR for detecting ^13^C-^13^C long-range correlations in proteins and polysaccharides. J. Biomol. NMR 61, 97–107 (2015).

71. Sauvée, C., et al. Highly efficient, water-soluble polarizing agents for dynamic nuclear polarization at high frequency. Angew. Chem. Int. Ed. 125, 11058-11061 (2013).

72 Dubroca, T. et al. A quasi-optical and corrugated waveguide microwave transmission system for simultaneous dynamic nuclear polarization NMR on two separate 14.1 T spectrometers. J. Magn. Reson. 289, 35–44 (2018).

73 De Paëpe, G., Lewandowski, J. R., Loquet, A., Böckmann, A. & Griffin, R. G. Proton assisted recoupling and protein structure determination. J Chem Phys 129, 12B615 (2008).

74 Pauli, J., Baldus, M., van Rossum, B., de Groot, H. & Oschkinat, H. Backbone and Side-Chain 13C and 15N Signal Assignments of the α-Spectrin SH3 Domain by Magic Angle Spinning Solid-State NMR at 17.6 Tesla. ChemBioChem 2, 272–281 (2001).

75 Jo, S., Kim, T., Iyer, V. G. & W., I. CHARMM-GUI: A web-based graphical user interface for CHARMM. J. Comput. Chem. 29, 1859–1865 (2008).

76 Jo, S., Song, K. C., Desaire, H., MacKerell, A. D. & Im, W. Glycan reader: Automated sugar identification and simulation preparation for carbohydrates and glycoproteins. J. Comput. Chem. 32, 3135–3141 (2011).

77. Malaspina, D. C. & Faraudo, J. Chitin builder (v1.0). Zenodo, https://doi.org/10.5281/zenodo.3274726 (2019).

78 Humphrey, W., Dalke, A. & Schulten, K. VMD: Visual molecular dynamics. J. Mol. Graph. 14, 33–38 (1996).

79 Rinaudo, M. Chitin and chitosan: Properties and applications. Prog. Polym. Sci. 31, 603–632 (2006).

80 Guvench, O., Hatcher, E., Venable, R. M., Pastor, R. W. & MacKerell, A. D. CHARMM Additive All-Atom Force Field for Glycosidic Linkages between Hexopyranoses. J. Chem. Theory Comput. 5, 2353–2370 (2009).

81 Guvench, O. et al. Additive empirical force field for hexopyranose monosaccharides. J. Comput. Chem. 29, 2543–2564 (2008).

82 Jorgensen, W. L., Chandrasekhar, J., Madura, J. D., Impey, R. W. & Klein, M. Comparison of simple potential functions for simulating liquid wate. J. Chem. Phys. 79, 926–935 (1983).

83 Beglov, D. & Roux, B. Finite representation of an infinite bulk system: Solvent boundary potential for computer simulations. J. Chem. Phys. 100, 9050–9063 (1994).

84 Essmann, U. et al. A smooth particle mesh Ewald method. J. Chem. Phys. 103, 8577–8593 (1995).

85 Miyamoto, S. & Kollman, P. A. Settle: An analytical version of the SHAKE and RATTLE algorithm for rigid water models. J. Comput. Chem. 13, 952–962 (1992).

86 Paterlini, M. G. & Ferguson, D. M. Constant temperature simulations using the Langevin equation with velocity Verlet integration. Chem. Phys. 236, 243–252 (1998).

87 Feller, S. E., Zhang, Y., Pastor, R. W. & Brooks, B. R. Constant pressure molecular dynamics simulation: The Langevin piston method. J. Chem. Phys. 103, 4613–4621 (1995).

88 Harris, C. R. et al. Array programming with NumPy. Nature 585, 357–362 (2020).

89 Fiorin, G., Klein, M. L. & Henin, J. Using collective variables to drive molecular dynamics simulations. Mol. Phys. 111, 3345–3362 (2013).

