## Supplementary Information for "Structural Remodeling of Fungal Cell Wall Promotes Resistance to Echinocandins"

### Table of Content


Supplementary Figure 1. Replication of *A. fumigatus* samples under caspofungin treatment 3

Supplementary Figure 2. Structure of β-glucans from chemical analysis 4

Supplementary Figure 3. Mobile domains containing heteropolysaccharides of GAG and GM 5

Supplementary Figure 4. Protein components retained after caspofungin treatment 6

Supplementary Figure 5. $^{13}$C-$T_1$ relaxation data of 3-day-old *A. fumigatus* cell walls 7

Supplementary Figure 6. DNP detection of proteins and lipids 8

Supplementary Figure 7. Atomic models used for all-atom MD modeling 9

Supplementary Table 1. Average cell wall thickness of comparable-sized hyphae 10

Supplementary Table 2. Molar composition of rigid polysaccharides 11

Supplementary Table 3. Chemical analysis using GC-MS 12

Supplementary Table 4. Molar composition of mobile polysaccharides 13

Supplementary Table 5. Glucan comparison between chemical analysis and ssNMR 14

Supplementary Table 6. Relative ratios between polysaccharides, proteins, and lipids 15

Supplementary Table 7. Water-edited intensities of polysaccharide carbon sites 16

Supplementary Table 8. $^{13}$C-$T_1$ relaxation times of polysaccharides in *A. fumigatus* cell walls 17

Supplementary Table 9. Solid-state NMR experimental parameters 18

Supplementary Table 10. Chemical shifts of biomolecules at ambient temperature 19

Supplementary Table 11. Chemical shifts of polysaccharides at DNP condition. 20

Supplementary Table 12. Intermolecular correlation in 3-day-old cell walls 21

Supplementary References 23

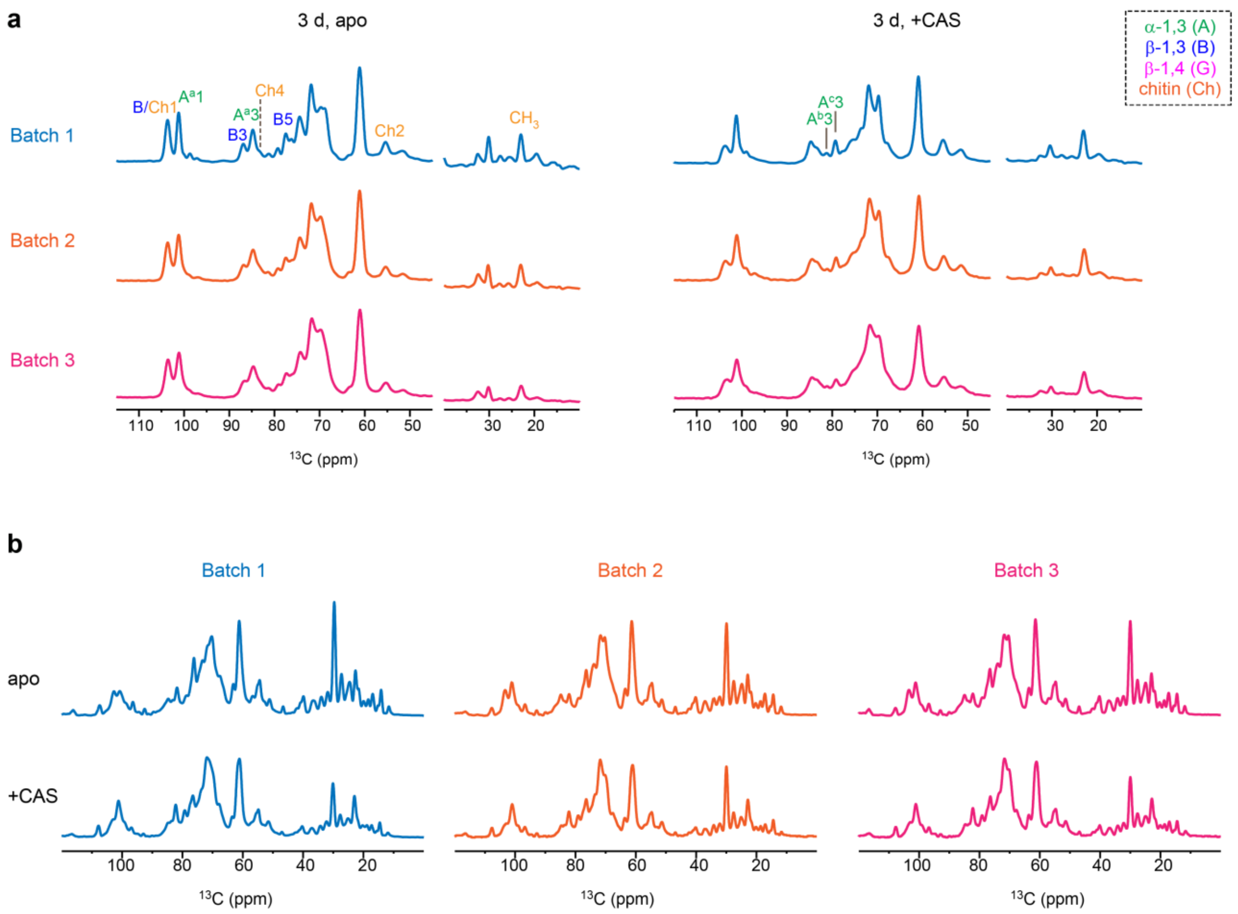


**Supplementary Figure 1. Replication of *A. fumigatus* samples under caspofungin treatment. a,** 1D $^{13}$C CP spectra of 3-day-old sample without drug (left) and with drug (right). **b**, Quantitative $^{13}$C DP spectra (with a recycle delay of 35 s) of 3-day-old without drug (top) and with drug (bottom). The carbohydrate region is highly replicable. The only noticeable change occurs at the 32-ppm peak, which is the lipid acyl chain ($CH_2$).

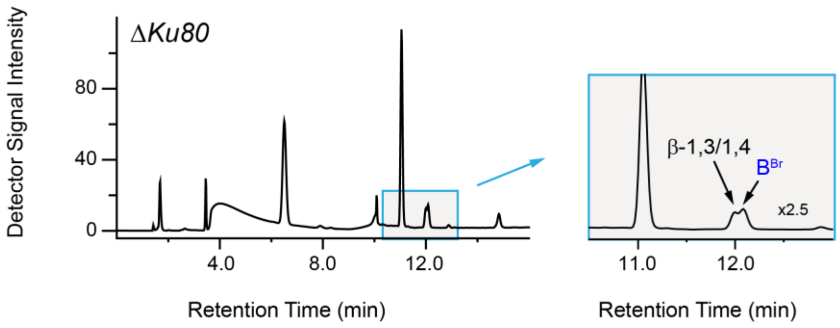


**Supplementary Figure 2. Structure of β-glucans from chemical analysis.** Data obtained from the Δ*akuB*$^{KU80}$ strain, which of is another widely used model strain of *A. fumigatus*, allow us to validate the results obtained on the wild-type strain used in this study. The right column shows the zoom-in regions where the different types of β-linkages could be resolved.

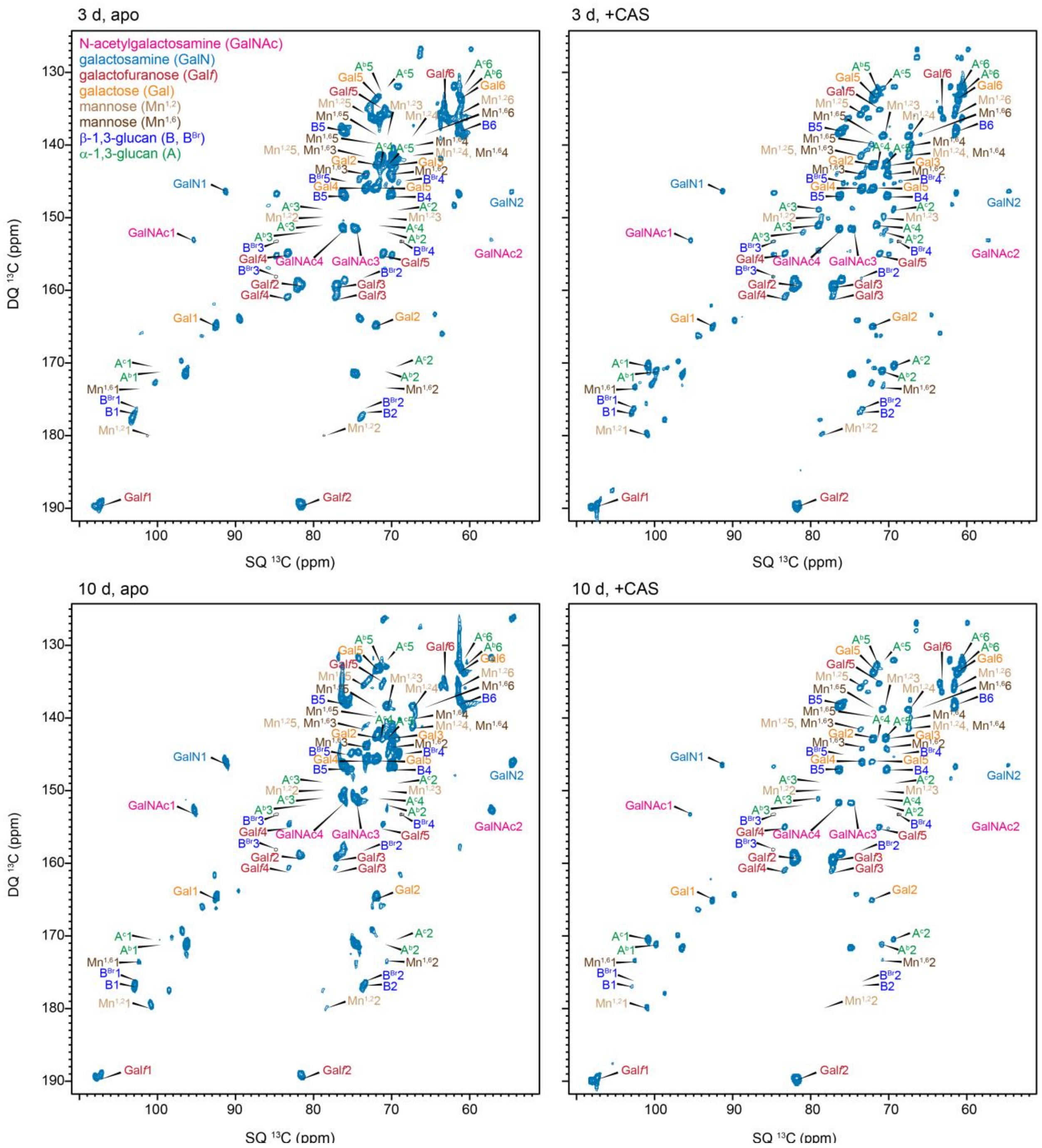


**Supplementary Figure 3. Mobile domains containing heteropolysaccharides of GAG and GM.** GAG contains N-acetylgalactosamine (GalNAc), galactosamine (GalN), and galactose (Gal) residues. GM contains galactofuranose (Gal*f*) and mannose residues with 1,2- and 1,6-linkages ($Mn^{1,2}$ and $Mn^{1,6}$). In both polysaccharides, sugar residues are arranged in no particular order. $B^{Br}$ indicates branching point in the β-(1,3)-linked backbone. Their chemical shifts are consistent in all four samples. The $^{13}C$ DP refocused J-INADEQUATE spectra detect the $^{13}C$ through-bond connectivity for mobile polysaccharides. All spectra were measured on 800 MHz NMR spectrometer at 12 kHz MAS.

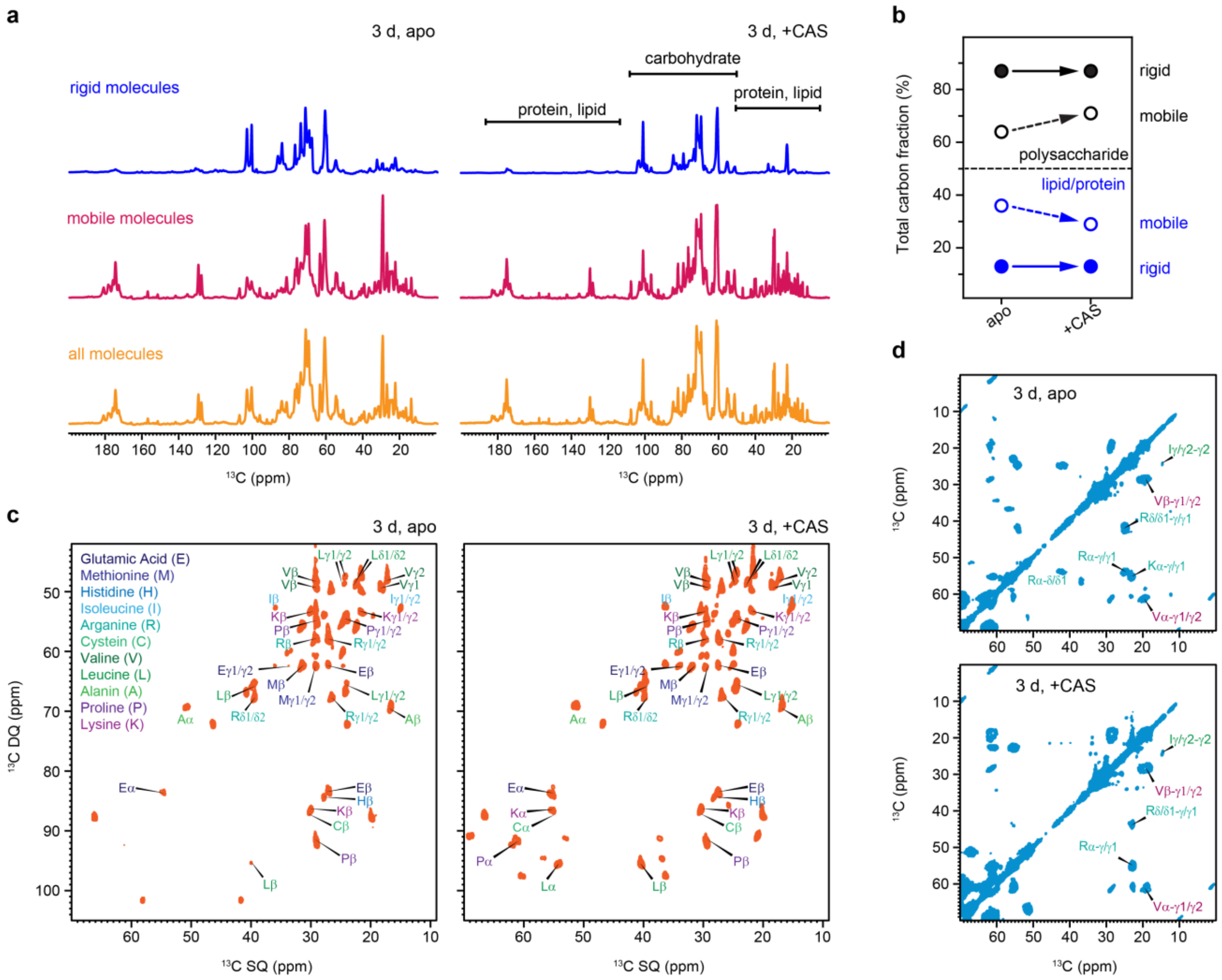

**Supplementary Figure 4. Protein components are retained after caspofungin treatment**. **a**, 1D $^{13}C$ NMR spectra measured using different pulse sequences: 1D $^{13}C$ CP for detecting rigid molecules, 2 s DP for detecting mobile molecules, and 30 s DP for quantitatively detecting all molecules. The spectra were collected from a 3-day-old sample both without (left) and with (right) caspofungin treatment. Specific regions corresponding to polysaccharides and protein/lipids were highlighted. **b**, Quantification of the total carbon fraction (%) of polysaccharides (black) and lipid/protein (blue) content within the sample. The sample fractions within mobile and rigid phases are indicated by open and filled circles, respectively. The alteration in the protein/lipid to polysaccharide ratio was observed exclusively in the mobile phase, with a minor change of less than 10%. **c**, 2D $^{13}C$ DP refocused J-INADEQUATE spectra showing signals of mobile proteins, with highly overlapping signals that indicate similar protein structures. **d**, 2D $^{13}C$-$^{13}C$ 53 ms CORD correlation spectra showing the rigid proteins. All spectra were measured on 800 MHz NMR spectrometer at 12 kHz MAS. Source data are provided as a Source Data file.

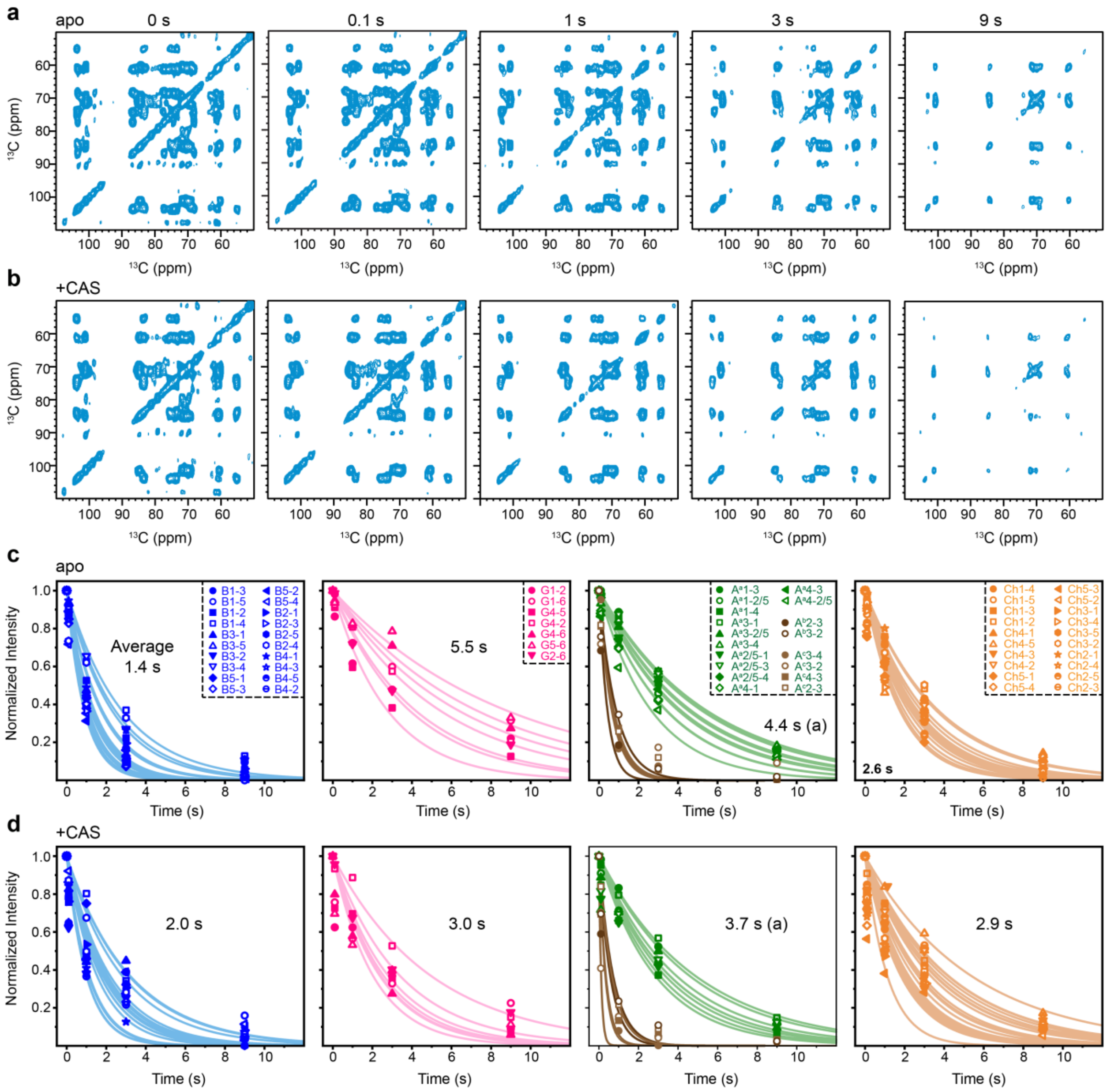


**Supplementary Figure 5. $^{13}$C-T$_1$ relaxation data of 3-day-old *A. fumigatus* cell walls.** Two arrays of 2D $^{13}$C-$^{13}$C spectra are shown for **a**, the *apo* sample and **b**, caspofungin-treated cell walls. From left to right, the z-filter duration increases (0 s, 0.1 s, 1 s, 3 s, and 9 s), and the intensities decrease. The intensity decay was plotted as a function of z-filter time for each carbon site. The relaxation decay curves are shown separately for **c,** the apo sample and **d,** the caspofungin-treated sample. The average $^{13}$C-T$_1$ time constants of polysaccharides are labeled. For α-1,3-glucan, the type-a subform is shown as green while the type-b and c forms in brown, with significantly faster relaxation.

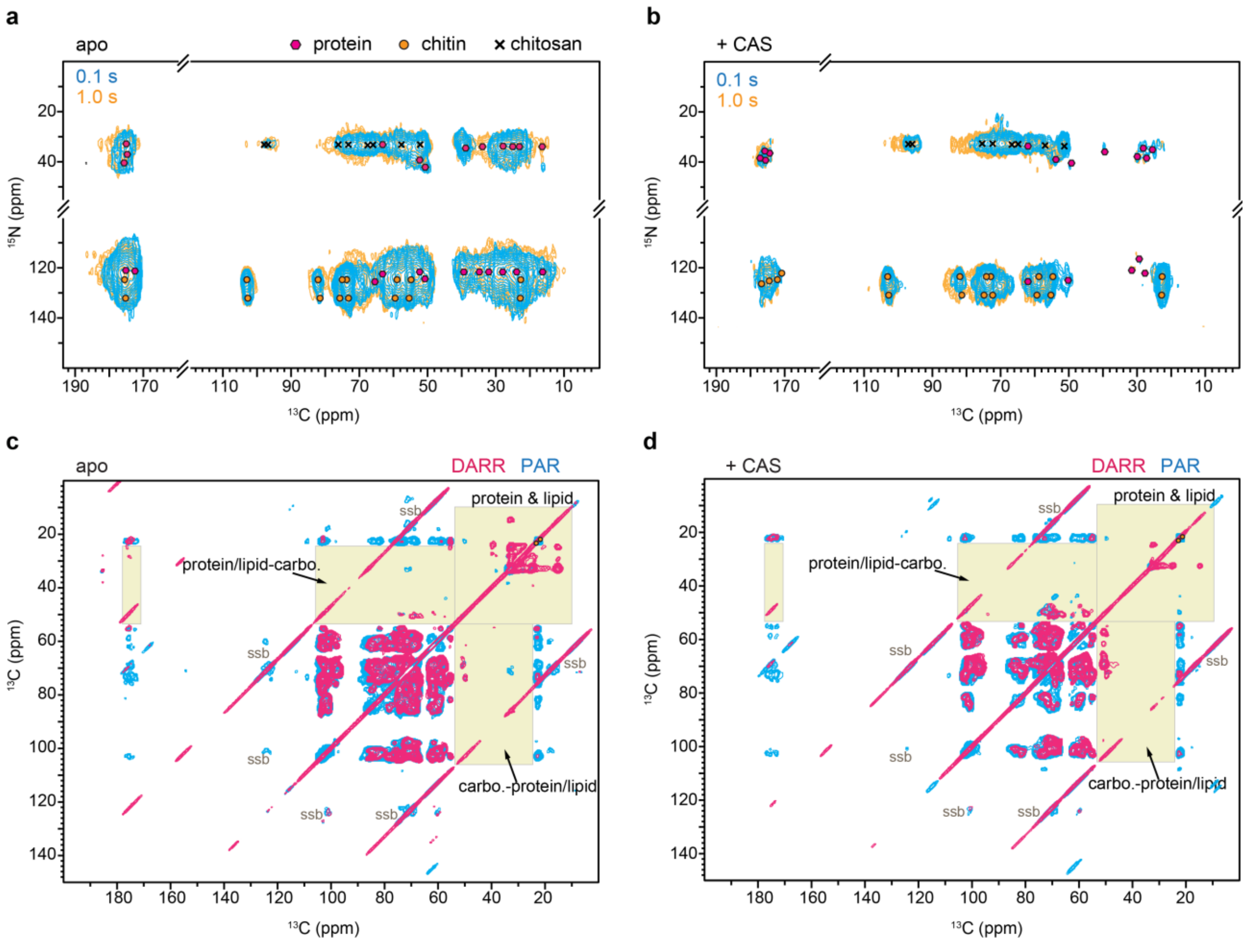


**Supplementary Figure 6. DNP detection of proteins and lipids in 3-day-old *A. fumigatus*.** 2D $^{15}$N-$^{13}$C correlation spectra of **a**, apo and **b**, drug-treated 3-day-old *A. fumigatus* samples exhibit signals from chitin NH (~125-135 ppm), chitosan $NH_2$ (~33 ppm), and some protein amide (~35-45 ppm). Overlay of spectra with 0.1-s and 1.0-s mixing times detect intra and inter-molecular cross peaks. 2D $^{13}$C-$^{13}$C correlation spectra of **c**, apo and **d**, drug-treated samples did not show clear cross peaks between protein/lipid and polysaccharides.

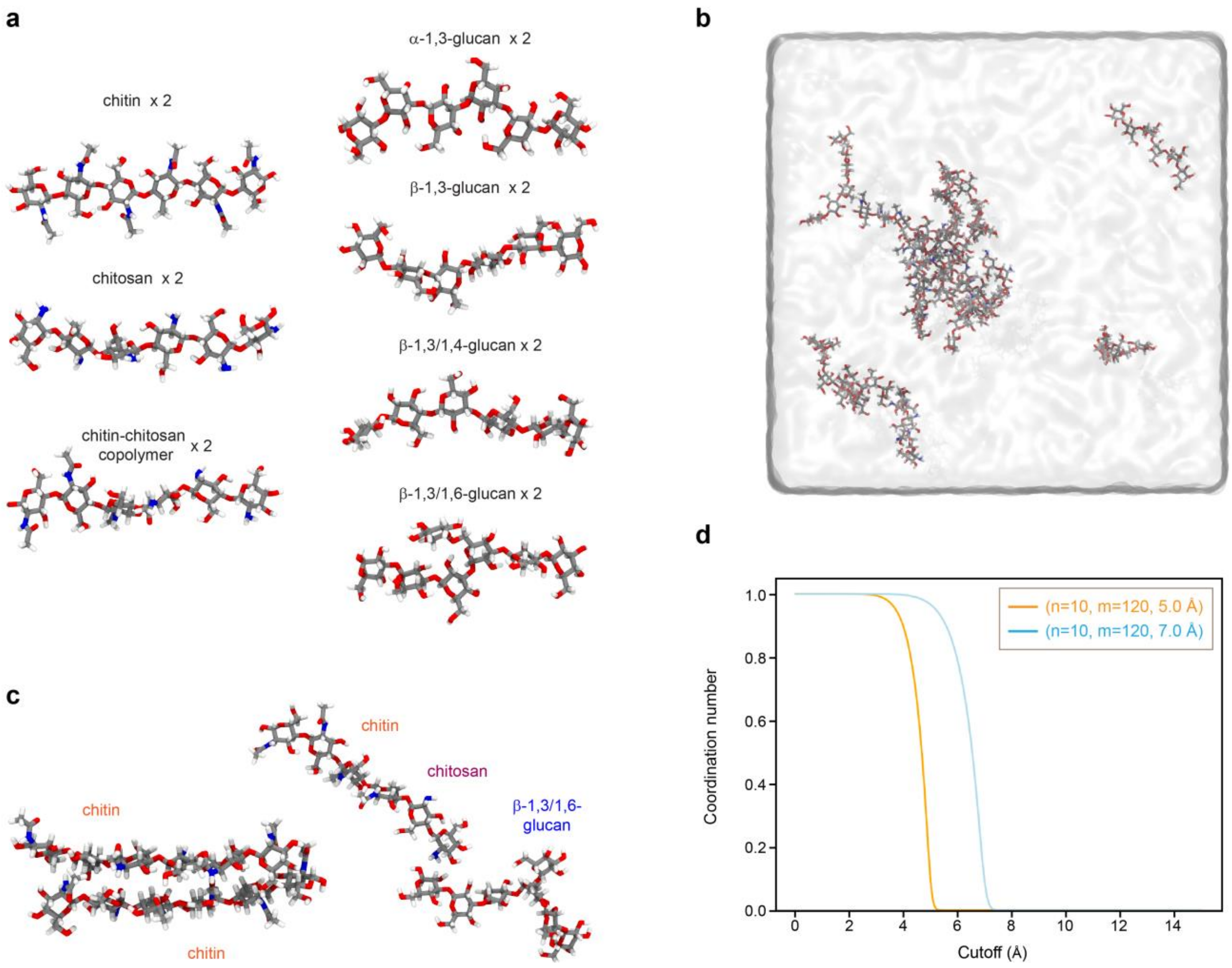


**Supplementary Figure 7. Atomic models used for all-atom MD modeling. a,** Atomic model of individual components included in the fungal cell wall molecular system. In total 14 molecules were included in the modeling, with two copies for each of the seven types of polysaccharides: chitin, chitosan, chitin-chitosan copolymer, α-1,3-glucan, linear β-1,3-glucan, branched β-1,3/1,6-glucan, and terminal β-1,3/1,4-glucan. **b,** Assembly of the molecular system solvated with water. Molecular structures for different polymers using VMD 1.9.4. The atomistic model consists of 120, 972 atoms after adding solvent and ions. **c,** Short-range interactions between two chitin chains, and between chitin-chitosan copolymer and β-1,3-glucan observed during 1 μs all-atom MD. **d,** Coordination number for short-range and long-range cutoff distance of 5.0 and 7.0 Å respectively, with the exponents n =10 and m = 120. See Methods for mathematical expression.

**Supplementary Table 1. Average cell wall thickness of comparable-sized hyphae**. Results are described as the averages and standard deviation of ~200 measurements of 10 individual cells in each sample. Error bars are standard deviation of each reading. A source data file is provided to document each single reading.

| Sample | 3 d, apo | 3 d, +drug | 10 d, apo | 10 d, +drug |
|---|---|---|---|---|
| Average cell wall thickness (nm) * | 133 ± 24 | 182 ± 29 | 123 ± 26 | 185 ± 32 |

Statistical analysis was performed using a one-tailed, paired t-test at 95% confidence level (*, $p < 0.05$) compared to the control condition (3-days-old).

**Supplementary Table 2. Molar composition of polysaccharides in the rigid phase.** The numbers are estimated using integrals (volume) of cross peaks in 2D 53-ms CORD $^{13}$C-$^{13}$C spectra. The average integrals of cross-peaks of each polysaccharide are shown. Error bars are standard errors of the peak integrals.

| **3 d, apo** | | | | | | | |
|---|---|---|---|---|---|---|---|
| β-glucans | | α-1,3-glucan | | | chitin | chitosan | |
| 38.7% | | 43.9% | | | | 1.8% | |
| β-1,3- | β-1,4 | a | b | c | | a | b |
| 35 ± 3 | 3.7 ± 0.9 | 39 ± 5 | 3 ± 1 | 1.9 ± 0.3 | 15 ± 2 | 0.8 ± 0.1 | 1.0 ± 0.1 |

| **3 d, +drug** | | | | | | | |
|---|---|---|---|---|---|---|---|
| β-glucans | | α-1,3-glucan | | | chitin | chitosan | |
| 3.7% | | 61% | | | | 12% | |
| β-1,3 | β-1,4 | a | b | c | | a | b |
| 1.2 ± 0.2 | 2.5 ± 0.4 | 27 ± 5 | 21 ± 4 | 13 ± 4 | 23 ± 2 | 6 ± 1 | 6 ± 3 |

| **10 d, apo** | | | | | | | |
|---|---|---|---|---|---|---|---|
| β-glucans | | α-1,3-glucan | | | chitin | chitosan | |
| 33% | | 58% | | | | | |
| β-1,3 | β-1,4 | a | b | c | | a | b |
| 16± 1 | 17 ± 3 | 36 ± 5 | 10.6 ± 0.7 | 8 ± 2 | 9 ± 1 | 2 ± 1 | 1.0 ± 0.1 |

| **10 d, +drug** | | | | | | | |
|---|---|---|---|---|---|---|---|
| β-glucans | | α-1,3-glucan | | | chitin | chitosan | |
| 5% | | 67% | | | | | |
| β-1,3 | β-1,4 | a | b | c | | a | b |
| 0.7 ± 0.1 | 3.3 ± 0.4 | 33 ± 5 | 16 ± 3 | 16 ± 3 | 20 ± 2 | 7 ± 1 | 5 ± 2 |

The area of the following well-resolved cross peaks 53-ms CORD spectra are used:
β-1,3: the average of C1-C2/3/4/5, C2-C4, C3-C2/4/5/6, and C5-C2/4/6.
β-1,4: the average of C1-6, C2-6, C4-6, and C5-6.
α-1,3-glucan (type a): the average of C1-C2/3/5/6 and C3-2/5/6.
α -1,3-glucan (type b): the average of C1-C2 and C3-2.
α-1,3-glucan (type c): the average of C1-C3 and C3-2/4/6.
Chitin: the average of C1-2/4/5, C3/5-2, C4-2/3/5, and C6-2.
Chitosan (type a and b): the average of C1-C2 and C3-C2.

**Supplementary Table 3. Chemical analysis using GC-MS.** The same samples prepared using minimum medium and analyzed by ssNMR were subjected to chemical analysis. Chemical analysis was done using GC-MS coupled with enzymatic degradation. Alkali-insoluble (AI), alkali-soluble (AS), Undetected (UD).

| Component | 3 d, apo | | 3 d, +drug | |
|---|---|---|---|---|
| | AI | AS | AI | AS |
| Glucose | 49.5 | 60 | 23 | 51 |
| Galactose | 18 | 22 | 17 | 28 |
| Mannan | 5.5 | 1 | 4 | 1 |
| GlcNAc | 23 | UD | 47 | UD |
| GalNAc | 4 | 17 | 9 | 20 |

**Supplementary Table 4. Molar composition of mobile polysaccharides in cell walls.** The numbers are estimated using integrals (volume) of spin pair peaks in 2D J-INADEQUATE $^{13}C$-$^{13}C$ spectra. The average integrals of cross-peaks of each polysaccharide are shown. Error bars are standard errors of the peak integrals. Undetected: UD.

| **3 d, apo** | | | | | | | | | | | |
|---|---|---|---|---|---|---|---|---|---|---|---|
| β-1,3-1,6-glucans | | α-1,3-glucan | | Galactosaminogalactan (GAG) | | | Galactomannan (GM) | | | Chitosan | |
| 26% | | 4% | | 48% | | | 20% | | | 2% | |
| β-1,3 | β-1,3 (Br) | b | c | Gal | GalN | GalNAc | $Mn^{1,2}$ | $Mn^{1,6}$ | Gal*f* | a | b |
| 24 ± 4 | 1.8 ± 0.4 | 3 ± 2 | 1 ± 1 | 33 ± 4 | 9 ± 1 | 6.2 ± 0.7 | 3 ± 3 | 2.2 ± 0.8 | 15 ± 2 | UD | 1.7 ± 0.3 |

| **3 d, +drug** | | | | | | | | | | | |
|---|---|---|---|---|---|---|---|---|---|---|---|
| β-1,3-1,6-glucans | | α-1,3-glucan | | Galactosaminogalactan (GAG) | | | Galactomannan (GM) | | | Chitosan | |
| 10% | | 33% | | 24% | | | 23% | | | 11% | |
| β-1,3 | β-1, 3 (Br) | b | c | Gal | GalN | GalNAc | $Mn^{1,2}$ | $Mn^{1,6}$ | Gal*f* | a | b |
| 7 ± 2 | 3.3 ± 0.5 | 12 ± 1 | 21 ± 2 | 15 ± 3 | 5 ± 1 | 3.4 ± 0.3 | 7 ± 2 | 5.6 ± 0.4 | 10 ± 1 | 6 ± 1 | 5 ± 1 |

| **10 d, apo** | | | | | | | | | | | |
|---|---|---|---|---|---|---|---|---|---|---|---|
| β-1,3-1,6-glucans | | α-1,3-glucan | | Galactosaminogalactan (GAG) | | | Galactomannan (GM) | | | Chitosan | |
| 13% | | 3% | | 67% | | | 15% | | | 1% | |
| β-1,3 | β-1, 3 (Br) | b | c | Gal | GalN | GalNAc | $Mn^{1,2}$ | $Mn^{1,6}$ | Gal*f* | a | b |
| 13.3 ± 0.8 | UD | 1.7 ± 0.6 | 1 ± 1 | 35 ± 4 | 19 ± 1 | 13 ± 1 | 6.3 ± 0.3 | 5 ± 2 | 4 ± 1 | 0.25 ± 0.02 | 0.9 ± 0.1 |

| **10 d, +drug** | | | | | | | | | | | |
|---|---|---|---|---|---|---|---|---|---|---|---|
| β-1,3-1,6-glucans | | α-1,3-glucan | | Galactosaminogalactan (GAG) | | | Galactomannan (GM) | | | Chitosan | |
| 7% | | 28% | | 25% | | | 35% | | | 7% | |
| β-1,3 | β-1, 3 (Br) | b | c | Gal | GalN | GalNAc | $Mn^{1,2}$ | $Mn^{1,6}$ | Gal*f* | a | b |
| 6 ± 2 | 0.66 ± 0.05 | 11.7 ± 0.9 | 16 ± 5 | 13 ± 2 | 7 ± 1 | 4.6 ± 0.8 | 9 ± 1 | 9.3 ± 0.9 | 16 ± 4 | 5 ± 2 | 2.0 ± 0.6 |

The area of the following well-resolved cross peaks 53-ms CORD spectra are used: the average of C1 and C2 spin pair for β-1,3, β-1,3 (Br), α-1,3 (b and c), Gal, GalN, GalNAc, $Mn^{1,2}$, and $Mn^{1,6}$, the average of C4 and C3 spin pair for Gal*f*.

**Supplementary Table 5. Glucan comparison between chemical analysis and ssNMR.** Alkali-insoluble (AI), alkali-soluble (AS), the fraction by whole cell wall ($F_{cw}$), glucan in each fraction ($F_{Glc}$), β-glucan in each fraction ($F_{\beta}$), α-glucan in each fraction ($F_{\alpha}$), polysaccharide percentage in whole cell wall using GC-MS ($poly_{GC\text{-}MS}$), polysaccharide percentage in rigid part of the cell wall using solid-state NMR ($poly_{ssNMR}$). $F_{cw}$, $F_{Glc}$, $F_{\beta}$, and $F_{\alpha}$ obtained by GC-MS and enzymatic degradation.

| Sample | $F_{cw}$ | | $F_{Glc}$ | $F_{\beta}$ | $F_{\alpha}$ | $poly_{GC\text{-}MS}$ | | $poly_{ssNMR}$ | |
|---|---|---|---|---|---|---|---|---|---|
| | | | | | | α-1,3 | β-1,3/1,4 | α-1,3 | β-1,3/1,4 |
| 3 d, apo | AI | 50.73 | 49.5 | 94.44 | 5.56 | 54 | 46 | 53 | 47 |
| | AS | 49.27 | 60 | 5.25 | 94.75 | | | | |
| 3 d, +drug | AI | 42.05 | 22 | 83.71 | 16.29 | 79 | 21 | 94 | 6 |
| | AS | 57.95 | 51 | 1.15 | 98.85 | | | | |

**Supplementary Table 6. Relative ratios between polysaccharides, proteins, and lipids.** The ratios denote the fractions of carbons distributed in the three types of molecules, which are quantified using the areas of their corresponding spectral regions in the 1D spectra. The lipid/protein-to-polysaccharide ratios were similar in the rigid phase but changed in the mobile domains.

| Type of experiment | | Component | 3 d, apo | 3 d, +drug | 10 d, apo | 10 d, +drug |
|---|---|---|---|---|---|---|
| 1D CP | Rigid | Polysaccharides | 0.87 | 0.87 | 0.88 | 0.85 |
| | | Lipids + proteins | 0.13 | 0.13 | 0.12 | 0.15 |
| Difference (30s - 2s) | | Polysaccharides | 0.86 | 0.87 | 0.88 | 0.87 |
| | | Lipids + proteins | 0.14 | 0.13 | 0.12 | 0.13 |
| 1D DP (2 s) | Mobile | Polysaccharides | 0.64 | 0.71 | 0.77 | 0.74 |
| | | Lipids + proteins | 0.36 | 0.29 | 0.23 | 0.26 |
| 1D DP (30 s) | All | Polysaccharides | 0.70 | 0.75 | 0.80 | 0.78 |
| | | Lipids + proteins | 0.30 | 0.25 | 0.20 | 0.22 |

The integral of the following areas in 1D CP, 1D DP (2 s), and 1D DP (30 s) spectra are used: 52-92 ppm and 98-108 ppm for polysaccharides; 2-52 ppm for lipids and proteins.

**Supplementary Table 7. Water-edited intensities of polysaccharide carbon sites.** The intensity ratios are obtained by comparing the peak intensities in water-edited and control 2D spectra. The average values for each molecule in each sample are highlighted in bold. Error bars are standard deviations propagated from NMR signal-to-noise ratios. The error margin is typically below 10% of the reported values. Only those error bars above 10% of the reported values are shown.

| Cross peaks | 3 d, apo | 3 d, drug | Cross peaks | 3 d, apo | 3 d, drug |
|---|---|---|---|---|---|
| **Average** | **0.68** | **0.38** | **Average** | **0.37** | **0.27** |
| B1-2 | 0.63 | 0.31 | $A^a$1-2/5 | 0.36 | 0.30 |
| B1-3 | 0.78 | 0.6±0.5 | $A^a$1-3 | 0.34 | 0.28 |
| B1-4 | 0.74 | 0.47±0.04 | $A^a$1-4 | 0.41 | 0.33 |
| B1-5 | 0.76 | - | $A^a$1-6 | 0.38 | 0.27 |
| B1-6 | 0.71 | 0.38 | $A^a$2/5-1 | 0.39 | 0.32 |
| B2-1 | 0.63 | 0.31 | $A^a$2/5-3 | 0.37 | 0.27 |
| B2-3 | 0.76 | - | $A^a$2/5-4 | 0.43 | - |
| B2-4 | 0.74 | 0.50 | $A^a$2/5-6 | 0.39 | 0.31 |
| B2-5 | 0.66 | 0.26 | $A^a$3-1 | 0.33 | 0.27 |
| B2-6 | 0.63 | 0.28 | $A^a$3-2/5 | 0.35 | 0.27 |
| B3-1 | 0.80 | - | $A^a$3-4 | 0.42 | 0.24 |
| B3-2 | 0.78 | 0.4±0.3 | $A^a$3-6 | 0.37 | 0.30 |
| B3-4 | 0.71 | 0.45 | $A^a$4-1 | 0.36 | 0.25 |
| B3-5 | 0.84 | - | $A^a$4-2/5 | 0.35 | 0.21 |
| B3-6 | 0.66 | 0.22±0.08 | $A^a$4-3 | 0.35 | 0.23 |
| B4-1 | 0.73 | - | $A^a$4-6 | 0.39 | 0.24 |
| B4-2 | 0.63 | 0.49±0.02 | **Average** | **0.47** | **0.34** |
| B4-3 | 0.60 | - | $A^b$1-2 | 0.46 | 0.28 |
| B4-5 | 0.62 | 0.49±0.02 | $A^b$3-2 | 0.57 | 0.33 |
| B4-6 | 0.48 | 0.28 | $A^b$2-1 | 0.42 | 0.42 |
| B5-1 | 0.68 | 0.5±0.1 | $A^b$2-3 | 0.45 | 0.32 |
| B5-2 | 0.57 | 0.26 | **Average** | **0.33** | **0.30** |
| B5-3 | 0.74 | - | $A^c$1-3 | 0.43 | 0.31 |
| B5-4 | 0.58 | 0.35 | $A^c$3-1 | 0.2±0.1 | 0.34 |
| B5-6 | 0.64 | 0.38 | $A^c$3-4 | 0.28 | 0.29 |
| **Average** | **0.41** | **0.25** | $A^c$3-2 | 0.24 | 0.28 |
| Ch1-2 | 0.43 | 0.23 | $A^c$4-3 | 0.38 | 0.28 |
| Ch1-3 | 0.57 | 0.23 | $A^c$2-3 | 0.42 | 0.31 |
| Ch1-4 | 0.43 | 0.20 | **Average** | **0.81** | **0.46** |
| Ch1-5 | 0.40 | 0.24 | G4-6 | 0.9±0.2 | - |
| Ch1-6 | 0.71 | 0.38 | G1-6 | 0.75 | 0.5±0.3 |
| Ch2-1 | 0.34 | 0.20 | G2-6 | - | 0.38 |
| Ch2-3 | 0.33 | 0.25 | | | |
| Ch2-4 | 0.28±0.03 | 0.31 | | | |
| Ch2-5 | 0.20 | 0.29 | | | |
| Ch2-6 | 0.36±0.04 | 0.26 | | | |
| Ch3-1 | 0.49 | 0.24 | | | |
| Ch3-2 | 0.32 | 0.25 | | | |
| Ch3-4 | 0.41 | 0.35 | | | |
| Ch3-5 | 0.43 | 0.23 | | | |
| Ch3-6 | 0.43 | 0.32 | | | |
| Ch4-1 | 0.44 | 0.26 | | | |
| Ch4-2 | 0.36 | 0.18 | | | |
| Ch4-3 | 0.40 | 0.28 | | | |
| Ch4-5 | 0.35 | 0.16 | | | |
| Ch4-6 | 0.42 | 0.26 | | | |
| Ch5-1 | 0.61 | 0.29 | | | |
| Ch5-2 | 0.30 | 0.21 | | | |
| Ch5-3 | 0.32 | 0.25 | | | |
| Ch5-4 | 0.35 | 0.23 | | | |
| Ch5-6 | 0.52 | 0.24 | | | |

**Supplementary Table 8. $^{13}$C-$T_1$ relaxation times of polysaccharides in *A. fumigatus* cell walls.** Data are shown for the 3-day-old sample, with and without treatment by caspofungin. The average values for each molecule in each sample are highlighted in bold. The data were measured using 2D $^{13}$C-$^{13}$C correlation experiments. The data are fit using single exponential equations: $I(t) = e^{-t/T_1}$. Error bars are standard deviations of the fit parameters.

| Cross peaks | 3 d apo | 3 d drug | Cross peaks | 3 d apo | 3 d drug |
|---|---|---|---|---|---|
| **Average** | **1.4** | **2.0** | **Average** | **4.4** | **3.7** |
| B1-2 | 1.7±0.2 | 1.7±0.5 | A$^{a}$1-2/5 | 4.0±0.3 | 3.6±0.3 |
| B1-3 | 1.06±0.07 | 1.9±0.5 | A$^{a}$1-3 | 5.4±0.3 | 4.6±0.2 |
| B1-4 | 2.8±0.5 | 3.2±0.7 | A$^{a}$1-4 | 4.7±0.2 | 2.9±0.2 |
| B1-5 | 1.10±0.04 | 3.2±0.7 | A$^{a}$2/5-1 | 4.5±0.3 | 3.1±0.7 |
| B2-1 | 1.4±0.2 | 2.0 ±0.5 | A$^{a}$2/5-3 | 5.2±0.3 | 3.6±0.7 |
| B2-3 | 1.0±0.1 | - | A$^{a}$2/5-4 | 3.4±0.2 | - |
| B2-4 | 2.5±0.7 | 1.8±0.3 | A$^{a}$3-1 | 5.3±0.2 | 4.9±0.7 |
| B2-5 | 1.21±0.06 | 2.0±0.6 | A$^{a}$3-2/5 | 4.6±0.5 | 3.9±0.5 |
| B3-1 | 1.40±0.09 | 2±1 | A$^{a}$3-4 | 5.4±0.3 | 3.4±0.9 |
| B3-2 | 1.2±0.1 | 1.1±0.5 | A$^{a}$4-1 | 3.4±0.4 | - |
| B3-4 | 2.3±0.6 | - | A$^{a}$4-2/5 | 2.7±0.4 | - |
| B3-5 | 1.26±0.06 | - | A$^{a}$4-3 | 4.2±0.6 | - |
| B4-1 | 1.8±0.3 | 1.2±0.1 | **Average** | **0.7** | **0.4** |
| B4-2 | 1.3±0.2 | 1.0±0.4 | A$^{b}$3-2 | 0.4±0.1 | 0.20±0.05 |
| B4-3 | 1.2±0.2 | - | A$^{b}$2-3 | 0.9±0.2 | 0.6±0.2 |
| B4-5 | 1.27±0.03 | - | **Average** | **0.7** | **0.4** |
| B5-1 | 1.06±0.03 | 2.8±0.5 | A$^{c}$3-4 | 0.7±0.1 | 0.31±0.03 |
| B5-2 | 0.90±0.06 | 2.1±0.6 | A$^{c}$3-2 | 0.7±0.2 | 0.11±0.02 |
| B5-3 | 1.1±0.1 | 1.5±0.2 | A$^{c}$4-3 | 0.75±0.08 | 0.35±0.08 |
| B5-4 | 1.0±0.3 | 1.7±0.3 | A$^{c}$2-3 | 0.7±0.1 | 0.66±0.08 |
| **Average** | **2.6** | **2.9** | **Average** | **5.5** | **3.0** |
| Ch1-2 | 3.7±0.3 | 3.6±0.3 | G1-2 | 3.9±0.9 | 3±1 |
| Ch1-3 | 3.0±0.1 | 3.5±0.5 | G1-6 | 6±1 | 2.4±0.8 |
| Ch1-4 | 3.6±0.1 | 2.8±0.6 | G2-6 | 4.2±0.5 | 3.4±0.5 |
| Ch1-5 | 1.8±0.2 | 2.9±0.6 | G4-2 | 5.4±0.4 | 5.0±0.5 |
| Ch2-1 | 2.4±0.2 | 2.7±0.6 | G4-5 | 2.8±0.4 | 2.8±0.7 |
| Ch2-3 | 2.6±0.5 | 4.2±0.7 | G4-6 | 7.3±0.7 | 2.0±0.4 |
| Ch2-4 | 3.0±0.3 | 2.1±0.8 | G5-6 | 9±1 | 2.4±0.9 |
| Ch2-5 | 1.8±0.3 | 3.3±0.5 | | | |
| Ch3-1 | 2.5±0.1 | 2.0±0.7 | | | |
| Ch3-2 | 3.6±0.5 | 2.5±0.6 | | | |
| Ch3-4 | 2.0±0.4 | 1.9±0.5 | | | |
| Ch3-5 | 2.2±0.3 | 1.8±0.6 | | | |
| Ch4-1 | 2.6±0.5 | 3.8±0.9 | | | |
| Ch4-2 | 2.5±0.4 | 4±1 | | | |
| Ch4-3 | 2.8±0.6 | 4±1 | | | |
| Ch4-5 | 2.3±0.7 | 5.4±0.8 | | | |
| Ch5-1 | 1.8±0.1 | 2.0±0.6 | | | |
| Ch5-2 | 2.3±0.5 | 2.2±0.6 | | | |
| Ch5-3 | 2.5±0.6 | 1.1±0.6 | | | |
| Ch5-4 | 2.7±0.7 | 1.8±0.7 | | | |

**Supplementary Table 9. Solid-state NMR experimental parameters for fungal cell wall characterization.** T = sample temperature; $B_0$ = magnetic field; $\nu_{MAS}$ = MAS frequency; ns = number of scans; $d_1$ = recycle delay between scans; $t_{1,\,max}$ = maximum $t_1$ evolution time (for indirect dimension); $t_{1,\,inc}$ = increment for $t_1$ (for indirect dimension) evolution time; $\tau_{dw}$ = dwell time during direct FID acquisition; $\tau_{acq}$ = maximum acquisition time during direct FID detection; $\tau_{XY}$ = cross-polarization contact time during CP from channel X to channel Y; $\nu_{1H,\,dec}$ = dipolar decoupling field strength. DNP experiments are marked with asterisks.

| **Experiment** | **NMR Parameters** | | | | | | | | | | | | | | | **Samples** |
|---|---|---|---|---|---|---|---|---|---|---|---|---|---|---|---|---|
| | T (K) | $B_0$ (T) | $\nu_{MAS}$ (kHz) | ns | d1 (s) | $t_{1,\,max}$ (ms) | $t_{1,\,inc}$ (µs) | $\tau_{dw}$ (µs) | $\tau_{acq}$ (ms) | $\tau_{HC}$ (ms) | $\tau_{HN}$ (ms) | $\tau_{NC}$ (ms) | $\tau_{SD}$ (ms) | $\tau_{mix}$ (ms) | $\nu_{1H\,dec}$ (kHz) | |
| 1D $^{13}C$ CP | 298 | 18.8 | 12 | 2048 | 2 | | | 7 | 17 | 1 | | | | | 83 | 3 d apo<br>3 d CAS<br>10 d apo<br>10 d CAS |
| 1D $^{13}C$ DP | 298 | 18.8 | 12 | 64-128 | 2 or 30 | | | 7 | 28 | | | | | | 83 | |
| 2D $^{13}C$-$^{13}C$ with CORD mixing | 298 | 18.8 | 12 | 16 | 1.7 | 7 | 26 | 7.5 | 18 | 1 | | | | 53 $\tau_{CORD}$ | 83 | |
| 2D $^{13}C$-$^{13}C$ DP J-INADEQUATE | 298 | 18.8 | 12 | 8 | 1.5 | 10 | 20 | 7.5 | 19 | | | | | | 83 | |
| 2D $^{15}N$-$^{13}C$ N(CA)CO with DARR mixing | 298 | 18.8 | 12 | 64 | 1.7 | 6 | 100 | 7.5 | 16 | | 0.6 | 5 | | 100 $\tau_{DARR}$ | 83 | |
| 2D $^{13}C$-$^{13}C$ water-edited | 298 | 9.4 | 10 | 128-256 | 1.6 | 5 | 70 | 10 | 14 | 1 | | | 0, 4 | 50 $\tau_{PDSD}$ | 83 | |
| Pseudo 3D $^{13}C$-$T_1$ | 298 | 9.4 | 10 | 80 | 1.6 | 5 | 110 | 10 | 16 | 1 | | | | 50 $\tau_{PDSD}$ | 83 | |
| * 2D $^{13}C$-$^{13}C$ with PDSD mixing | 92 | 14.1 | 8 | 4 | 6.7 | 7 | 36 | 7.5 | 11 | 0.5 | | | | 100 $\tau_{PDSD}$ | 71 | 3 d apo<br>3 d CAS |
| * 2D $^{13}C$-$^{13}C$ with PAR mixing | 92 | 14.1 | 8 | 8 | 7.5 | 7 | 34 | 7.8 | 8 | 0.5 | | | | 20 $\tau_{PAR}$ | 71 | |
| * 2D $^{15}N$-$^{13}C$ N(CA)CO with PDSD | 92 | 14.1 | 8 | 32 | 8 | 3 | 124 | 7.5 | 13 | | 1 | 4 | | 100 or 3000 $\tau_{PDSD}$ | 71 | |

**Supplementary Table 10. $^{13}$C and $^{15}$N chemical shifts of biomolecules in *A. fumigatus* cell walls at ambient temperature.** Superscripts are used to denote different allomorphs. Not applicable (/). Unidentified (-). Branched (Br). Reducing end (O).

| Carbohydrate | | C1 | C2 | C3 | C4 | C5 | C6 | CO | $CH_3$ | N | Experiment | References |
|---|---|---|---|---|---|---|---|---|---|---|---|---|
| α-1,3-glucan | a | 101 | 71.9 | 84.6 | 69.5 | 71.7 | 60.5 | / | / | / | $^{13}$C-$^{13}$C CORD | Bhanja *et al*.2014[1] |
| | b | 99.9 | 71.0 | 81.1 | 73.1 | 71.9 | 60.8 | / | / | / | $^{13}$C-$^{13}$C CORD, $^{13}$C DP J-INADEQUATE | |
| | c | 100.9 | 69.4 | 79.1 | 71.9 | 70.8 | 60.9 | / | / | / | | |
| β-1,3-glucan | | 103.6 | 74.4 | 86.4 | 68.7 | 77.1 | 61.3 | / | / | / | $^{13}$C-$^{13}$C CORD | Shim *et al*. 2007[2] Fairweather *et al*. 2009[3] Saito *et al*. 1979[4] |
| β-1,4-glucan | | 103.3 | 69.4 | 71.7 | 85.3 | 74.3 | 63.4 | / | / | / | | Kang *et al*. 2018[5] |
| β-1,3-glucan ($B^{Br}$) | | 103.2 | 73.9 | 85.2 | 69.1 | 76.2 | 69.7 | / | / | / | $^{13}$C DP J-INADEQUATE | Lowman et al. 2011[6] |
| chitin | | 103.6 | 55.5 | 72.9 | 83.0 | 75.7 | 60.9 | 174.8 | 22.6 | 123.6 | $^{13}$C-$^{13}$C CORD | Kono *et al*. 2004[7] |
| chitosan | a | 98.8 | 50.9 | 67.3 | 76.9 | 71.7 | 60.2 | / | / | 32.9 | $^{13}$C-$^{13}$C CORD, $^{13}$C DP J-INADEQUATE | Fernando et al. 2021[8] |
| | b | 96.6 | 51.6 | 66.6 | 76.6 | 71.7 | 61.1 | / | / | 32.9 | | |
| $Mn^{1,2}$ | | 101.7 | 79.1 | 71.3 | 68.3 | 74.3 | 62.3 | / | / | / | $^{13}$C DP J-INADEQUATE | Latgé *et al*.1994[9] Chakraborty *et al*. 2021[10] |
| $Mn\text{-}O^{1,2}$ | | 99.4 | 79.1 | 71.3 | 68.3 | 74.3 | 62.3 | / | / | / | | |
| $Mn^{1,6}$ | | 101 | 72.9 | 73.8 | 67.9 | 72.8 | 66.6 | / | / | / | | |
| Gal*f* | | 107.5 | 81.6 | 77.7 | 83.5 | 71.5 | 63.5 | / | / | / | | |
| Gal | | 93.2 | 72.2 | 70.7 | 73.5 | 72.5 | 60.9 | / | / | / | | Fontaine *et al*. 2011[11] |
| GalN | | 91.7 | 54.8 | 71.1 | 76.9 | - | - | / | - | - | | |
| GalNAc | | 95.7 | 57.5 | 75.2 | 81.1 | - | - | 175.2 | 22.7 | - | | |

| Amino Acids | Cα | Cβ | Cγ/γ1 | Cγ2 | Cδ/δ1 | | Amino Acids | Cα | Cβ | Cγ/γ1 | Cγ2 | Cδ/δ1 | References |
|---|---|---|---|---|---|---|---|---|---|---|---|---|---|
| Glutamic Acid (E) | 55.2 | 27.2 | 33.8 | | | | Valine (V) | | 29.2 | 18.4 | | | Fritzsching *et al*. 2013[12] |
| Methionine (M) | | 32.0 | 29.5 | | | | Leucine (L) | 54.1 | 40.1 | 24.6 | | 22.4 | |
| Histidine (H) | | 27.9 | | | | | Alanine (A) | 51.3 | 16.6 | | | | |
| Arginine (R) | | 29.5 | 26.8 | | 39.5 | | Proline (P) | 61.1 | 29.1 | 24.8 | | | |
| Cysteine (C) | 55.0 | 30.5 | | | | | Lysine (K) | 55.1 | 30.9 | 21.5 | | | |
| Isoleucine (I) | | 36.1 | | 15.0 | | | | | | | | | |

**Supplementary Table 11. Chemical shifts of polysaccharides at DNP condition.** Superscripts are used to denote different allomorphs. Not applicable (/). Unidentified (-). Ambiguous (_).

<table>
<tr><th colspan="2">Carbohydrate</th><th>C1</th><th>C2</th><th>C3</th><th>C4</th><th>C5</th><th>C6</th><th>CO</th><th>$CH_3$</th><th>N</th><th>Experiment</th><th>Sample</th></tr>
<tr><td colspan="2">α-1,3-glucan</td><td>101.1</td><td>71.4</td><td>84.9</td><td>69.9</td><td>71.3</td><td>60.2</td><td>/</td><td>/</td><td>/</td><td rowspan="8">$^{13}C$-$^{13}C$ PAR and PDSD</td><td rowspan="11">3 d, apo</td></tr>
<tr><td colspan="2">β-1,3-glucan</td><td>103</td><td>74.1</td><td>84.6</td><td>68.9</td><td>77.8</td><td>61.7</td><td>/</td><td>/</td><td>/</td></tr>
<tr><td colspan="2">β-1,4-glucan</td><td>102.9</td><td>69.0</td><td><u>72.6</u></td><td>84.6</td><td>74.0</td><td>63.1</td><td>/</td><td>/</td><td>/</td></tr>
<tr><td rowspan="5">chitin</td><td>a</td><td>102.7</td><td>55.6</td><td>73.0</td><td>81.2</td><td>75.1</td><td>58.3</td><td>176.6</td><td>22.4</td><td></td></tr>
<tr><td>b</td><td>103.0</td><td>55.6</td><td>72.9</td><td>82.0</td><td>73.6</td><td>58.8</td><td>175.3</td><td>22.7</td><td></td></tr>
<tr><td>c</td><td>103.1</td><td>55.0</td><td>72.5</td><td>81.3</td><td>75.0</td><td>59.3</td><td>173.8</td><td>22.7</td><td></td></tr>
<tr><td>d</td><td>102.2</td><td>55.9</td><td>73.1</td><td>82.0</td><td>72.7</td><td>59.1</td><td>172.6</td><td>22.5</td><td></td></tr>
<tr><td>e</td><td>103.2</td><td>55.2</td><td>72.5</td><td>81.1</td><td>74.2</td><td>59.0</td><td>171.7</td><td>22.6</td><td></td></tr>
<tr><td rowspan="2">chitosan</td><td>a</td><td>98</td><td rowspan="2">51.9</td><td>67.3</td><td rowspan="2">76.9</td><td rowspan="2">73.1/<br>69.8</td><td rowspan="2">57.3</td><td>/</td><td>/</td><td>33.2</td><td rowspan="2">$^{15}N$-$^{13}C$ N(CA)CX with 100 ms DARR</td></tr>
<tr><td>b</td><td>96.8</td><td>65.7</td><td>/</td><td>/</td><td>33.2</td></tr>
<tr><td colspan="2">α-1,3-glucan</td><td>101.4</td><td>70.1</td><td>84.1</td><td>69.7</td><td>71.2</td><td>60.4</td><td>/</td><td>/</td><td></td><td rowspan="6">$^{13}C$-$^{13}C$ PAR and PDSD</td><td rowspan="8">3 d, + drug</td></tr>
<tr><td colspan="2">β-1,4-glucan</td><td>102.9</td><td><u>69.2</u></td><td><u>72.6</u></td><td>84.6</td><td>74.0</td><td>63.1</td><td>/</td><td>/</td><td></td></tr>
<tr><td rowspan="4">chitin</td><td>a</td><td>102.3</td><td>54.9</td><td>72.7</td><td>81.6</td><td>72.7</td><td>58.3</td><td>176.6</td><td>22.3</td><td></td></tr>
<tr><td>b</td><td>102.8</td><td>55.0</td><td>72.5</td><td>81.6</td><td>73.7</td><td>58.4</td><td>175.3</td><td>22.5</td><td></td></tr>
<tr><td>c</td><td>102.5</td><td>55.1</td><td>72.5</td><td>81.3</td><td>73.6</td><td>59.3</td><td>173.0</td><td>22.0</td><td></td></tr>
<tr><td>d</td><td>102.6</td><td>55.2</td><td>72.8</td><td>81.6</td><td>73.7</td><td>59.7</td><td>171.3</td><td>21.2</td><td></td></tr>
<tr><td rowspan="2">chitosan</td><td>a</td><td>98.8</td><td rowspan="2">51.6</td><td>67.1</td><td rowspan="2">76.4</td><td rowspan="2">73.5/<br>69.8</td><td rowspan="2">57.8</td><td>/</td><td>/</td><td>33.0</td><td rowspan="2">$^{15}N$-$^{13}C$ N(CA)CX with 100 ms DARR</td></tr>
<tr><td>b</td><td>96.1</td><td>65.7</td><td>/</td><td>/</td><td>33.0</td></tr>
</table>

**Supplementary Table 12. Intermolecular interactions in 3-day-old *Aspergillus* cell walls.** DNP-enhanced 20-ms PAR and $^{15}N$-$^{13}C$ N(CA)CX spectra with 3 s PDSD mixing are used to determine the intermolecular interactions. Cross peaks observed in 0.1 s PDSD and NCACX with 0.1 s PDSD mixing represent strong correlations. Mixed intermolecular interactions are underlined.

| 3d, apo | | | | | | |
|---|---|---|---|---|---|---|
| Cross peak | ω1 (ppm) | ω2 (ppm) | 0.1 s PDSD | 20 ms PAR | NCACX (0.1 s) | NCACX (3 s) |
| α-1,3-glucan-chitin | | | | | | |
| ChMe-A1 | 22.4 | 101.0 | | x | | |
| ChMe-A2,5 | 21.8 | 71.3 | | x | | |
| ChMe-A6 | 22.6 | 60.3 | | x | | |
| A2,5-ChCO | 70.8 | 175.1 | | x | | |
| A1-ChCO | 101.3 | 174.4 | | x | | |
| Ch4-A1 | 81.5 | 101.2 | x | x | | |
| ChN$_H$-A1 | 126.9 | 100.7 | | | | x |
| Ch1-A2,5 | 103.3 | 71.0 | x | x | | |
| A6-ChMe | 60.3 | 22.4 | | x | | |
| A2,5-ChMe | 70.3 | 22.5 | x | x | | |
| A1-ChMe | 100.9 | 22.7 | | x | | |
| A1-Ch4 | 101.1 | 82.0 | x | x | | |
| A3-ChMe | 84.1 | 22.5 | | x | | |
| A1-Ch1 | 100.9 | 103.2 | | x | | |
| Ch4-A3 | 81.5 | 84.4 | | x | | |
| A2,5-Ch1 | 70.8 | 103.1 | x | x | | |
| ChMe-A3 | 22.5 | 84.5 | | x | | |
| ChN$_H$-A3 | 124.4 | 84.2 | | | x | x |
| A3-Ch4 | 83.4 | 81.1 | | x | | |
| Ch1-A1 | 102.9 | 100.9 | | x | | |
| Ch4'-A3 | 81.1 | 83.8 | | x | | |
| α-1,3-glucan-chitosan | | | | | | |
| Cs1-A1 | 97.3 | 101.7 | | x | | |
| CsN-A1 | 33.0 | 100.1 | | | | x |
| A1-Cs1 | 101.3 | 97.2 | x | x | | |
| α-1,3-glucan-β-1,3-glucan | | | | | | |
| B5-A1 | 78.1 | 101.1 | | x | | |
| A1-B5 | 100.7 | 78.3 | | x | | |
| A1-B6 | 100.9 | 62.1 | x | x | | |
| A1-B1 | 100.9 | 103.2 | | x | | |
| A2,5-B1 | 70.8 | 103.1 | x | x | | |
| B1-A1 | 102.9 | 100.9 | | x | | |
| β-1,3-glucan-chitin | | | | | | |

| 3d, +drug | | | | | | |
|---|---|---|---|---|---|---|
| Cross peak | ω1 (ppm) | ω2 (ppm) | 0.1 s PDSD | 20 ms PAR | NCACX (0.1 s) | NCACX (3 s) |
| α-1,3-glucan-chitin | | | | | | |
| ChMe-A1 | 22.1 | 100.7 | | x | | |
| ChMe-A2,5 | 22.0 | 71.4 | | x | | |
| ChMe-A6 | 22.0 | 60.9 | | x | | |
| A2,5-ChCO | 69.5 | 173.2 | | x | | |
| A1-ChCO | 101.0 | 175.2 | | x | | |
| Ch4-A1 | 81.4 | 100.7 | | x | | |
| ChN$_H$-A1 | 126.8 | 101.1 | | | | x |
| Ch1-A2,5 | 102.7 | 70.0 | | x | | |
| A6-ChMe | 60.3 | 22.4 | | x | | |
| A2,5-ChMe | 69.3 | 22.0 | x | x | | |
| A1-ChMe | 100.7 | 21.9 | | x | | |
| A3-ChMe | 83.6 | 22.5 | | x | | |
| A1-Ch1 | 100.8 | 102.9 | | x | | |
| Ch4-A3 | 81.7 | 83.7 | | x | | |
| A2,5-Ch1 | 69.5 | 102.8 | | x | | |
| ChMe-A3 | 22.1 | 83.8 | | x | | |
| ChN$_H$-A3 | 124.3 | 84.1 | | | x | x |
| A3-Ch4 | 83.5 | 82.0 | | x | | |
| Ch1-A1 | 102.8 | 100.7 | | x | | |
| Ch1-A3 | 102.8 | 84.1 | | x | | |
| A3-Ch1 | 84.1 | 102.5 | | x | | |
| A6-ChCO | 60.4 | 173.0 | | x | | |
| A6-ChCO' | 60.4 | 175.3 | | x | | |
| Ch2-A3 | 55.6 | 83.7 | | x | | |
| A3-Ch2 | 83.6 | 55.6 | x | x | | |
| α-1,3-glucan-chitosan | | | | | | |
| Cs1-A1 | 98.2 | 100.9 | | x | | |
| CsN-A1 | 32.8 | 101.2 | | | | x |
| A1-Cs1 | 101.0 | 98.3 | | x | | |
| CsN-A3 | 33.2 | 84.0 | | | | x |
| β-1,3/1,4-glucan-chitin | | | | | | |
| ChMe-G6 | 22.0 | 62.1 | | x | | |
| Ch6-G6 | 58.1 | 62.8 | | x | | |

| | | | | | | |
|---|---|---|---|---|---|---|
| ChMe-B4 | 22.1 | 68.8 | x | x | | |
| ChMe-B6 | 22.5 | 62.1 | | x | | |
| B6-ChCO | 61.8 | 175.3 | | x | | |
| B6-ChCO’ | 62.0 | 173.5 | | x | | |
| B5-Ch4 | 77.6 | 80.6 | | x | | |
| B4-Ch4 | 68.4 | 81.7 | | x | | |
| B6-Ch4 | 61.3 | 81.6 | | x | | |
| Ch6-B5 | 58.9 | 78.2 | | x | | |
| Ch6-B6 | 58.8 | 62.2 | | x | | |
| B6-ChMe | 61.9 | 22.5 | x | x | | |
| B4-ChMe | 67.7 | 22.9 | | x | | |
| B6-Ch6 | 62.2 | 59.0 | | x | | |
| B5-Ch6 | 78.1 | 59.0 | | x | | |
| Ch4-B6 | 81.7 | 62.2 | | x | | |
| Ch4-B4 | 81.8 | 68.2 | | x | | |
| Ch4-B5 | 80.8 | 77.7 | | x | | |
| B3-ChMe | 84.1 | 22.5 | | x | | |
| B3-Ch4 | 84.4 | 82.5 | | x | | |
| ChMe-B3 | 22.5 | 84.5 | | x | | |
| Ch4-B3 | 81.5 | 84.4 | | x | | |
| Ch4’-B3 | 81.1 | 83.8 | | x | | |
| $ChN_H$-B3 | 124.4 | 84.2 | | | x | x |
| chitin-chitin’/chitosan | | | | | | |
| ChMe-Me’ | 22.9 | 21.4 | | x | | |
| Ch2-2’ | 54.6 | 55.8 | | x | | |
| Ch2’-2 | 55.8 | 54.7 | | x | | |
| ChMe’-Me | 21.4 | 22.9 | | x | | |
| CsN-Ch4 | 32.3 | 81.2 | | | | x |
| CsN-Ch1 | 33.4 | 104.0 | | | | x |
| β-1,3-glucan-chitosan | | | | | | |
| CsN-B1 | 33.4 | 104.0 | | | | x |

| | | | | | | |
|---|---|---|---|---|---|---|
| $ChN_H$-G4 | 124.9 | 85.6 | | | | x |
| G6- Ch6 | 62.4 | 58.3 | | x | | |
| G6-ChMe | 58.1 | 62.8 | | x | | |
| chitin-chitin’/chitosan | | | | | | |
| ChMe-Me’ | 22.9 | 21.2 | | x | | |
| Ch2-2’ | 54.3 | 55.7 | | x | | |
| Ch2’-2 | 55.5 | 54.3 | | x | | |
| ChMe’-Me | 21.2 | 22.9 | | x | | |
| ChCO-CO’ | 20.9 | 176.4 | | x | | |
| ChCO’-CO | 22.4 | 170.5 | | x | | |
| CsN-Ch4 | 33.5 | 81.5 | | | | x |
| CsN-Ch1 | 31.7 | 102.2 | | | | x |
| Cs1-Ch1 | 96.5 | 102.2 | | x | | |
| Ch1-Cs1 | 102.6 | 97.1 | | x | | |

#### Supplementary References


1. Bhanja, S. K. *et al.* Water-insoluble glucans from the edible fungus Ramaria botrytis. *Bioact. Carbohydr. Diet. Fibre* **3**, 52-58 (2014).
2. Shim, J. H. *et al.* Antitumor Effect of Soluble β-1, 3-Glucan from Agrobacterium sp. R259 KCTC 1019. *J. Microbiol. Biotechnol.* **17**, 1513-1520 (2007).
3. Fairweather, J. K., Him, J. L. K., Heux, L., Driguez, H. & Bulone, V. Structural characterization by $^{13}$C-NMR spectroscopy of products synthesized in vitro by polysaccharide synthases using $^{13}$C-enriched glycosyl donors: application to a UDP-glucose:(1→ 3)-b-d-glucan synthase from blackberry (Rubus fruticosus). *Glycobiology* **14**, 775-781 (2004).
4. Saitô, H., Ohki, T. & Sasaki, T. A 13C-nuclear magnetic resonance study of polysaccharide gels. Molecular architecture in the gels consisting of fungal, branched (1→ 3)-b-D-glucans (lentinan and schizophyllan) as manifested by conformational changes induced by sodium hydroxide. *Carbohydr. Res.* **74**, 227-240 (1979).
5. Kang, X. *et al.* Molecular architecture of fungal cell walls revealed by solid-state NMR. *Nat. Commun.* **9**, 1-12 (2018).
6. Lowman, D. W. *et al.* New Insights into the Structure of (1→3,1→6)-β-D-Glucan Side Chains in the Candida glabrata Cell Wall. *PLoS One* **6**, e27614 (2011).
7. Kono, H., Numata, Y., Erata, T. & Takai, M. $^{13}$C and $^{1}$H resonance assignment of mercerized cellulose II by two-dimensional MAS NMR spectroscopies. *Macromolecules* **37**, 5310-5316 (2004).
8. Fernando, L. D. *et al.* Structural polymorphism of chitin and chitosan in fungal cell walls from solid-state NMR and principal component analysis. *Front. Mol. Biosci.*, 727053 (2021).
9. Latge, J. P. *et al.* Chemical and immunological characterization of the extracellular galactomannan of Aspergillus fumigatus. *Infect. Immun.* **62**, 5424-5433 (1994).
10. Chakraborty, A. *et al.* A molecular vision of fungal cell wall organization by functional genomics and solid-state NMR. *Nat. Commun.* **12**, 1-12 (2021).
11. Fontaine, T. *et al.* Galactosaminogalactan, a new immunosuppressive polysaccharide of Aspergillus fumigatus. *PLoS Pathog.* **7**, e1002372 (2011).
12. Fritzsching, K., Yang, Y., Schmidt-Rohr, K. & Hong, M., Practical use of chemical shift databases for protein solid-state NMR: 2D chemical shift maps and amino-acid assignment with secondary-structure information. *J. Biomol. NMR* **2013,** *56*, 155-167.
