## Supplementary material for "Structural Remodeling of Fungal Cell Wall Promotes Resistance to Echinocandins": Captions of Supplementary Movies

**Supplementary Movie 1: Fungal cell wall interactions.mp4**
Molecular simulation of the interactions of polysaccharides in *A. fumigatus* cell walls. The specific composition used in the simulation is shown in **Supplementary Figure 7**. The solvent box is shown as a transparent glass surface and represents the field of view (FOV). Different polysaccharides are represented by spheres colored based on the atom – carbon (gray), oxygen (red), nitrogen (blue), hydrogen (white). The polymers inside the solvent box (FOV) appear as bright spheres, and fade as they diffuse into neighboring periodic images in the movie.

**Supplementary Movie 2: Alphaglucan chitin chitosan interactions.mp4**
Interaction between chitin, chitosan and α-1,3-glucan observed in a microsecond-long molecular simulation. The solvent box is shown as a transparent glass surface and the FOV. The polymers are represented by spheres and colored based on the atom type – carbon (gray), oxygen (red), nitrogen (blue), hydrogen (white). The polymers inside the solvent box (FOV) appear as bright spheres, and fade as they diffuse into neighboring periodic images in the movie. During the course of the simulation, the polysaccharides make both short (<5Å) and long-range (<10Å) interactions with each other.

**Supplementary Movie 3: Betaglucan chitin chitosan interactions.mp4**
Interaction between chitin, chitosan and β-glucan polymers observed in a microsecond-long molecular simulation. The solvent box is shown as a transparent glass surface and the FOV. The polymers are represented by spheres and colored based on the atom – carbon (gray), oxygen (red), nitrogen (blue), hydrogen (white). The polymers inside the solvent box (FOV) appear as bright spheres, and fade as they diffuse into neighboring periodic images in the movie. During the course of the simulation, the polysaccharides make both short (<5Å) and long-range (<10Å) interactions with each other.

**Supplementary Movie 4: Chitin chitosan interactions.mp4**
Interaction between chitin and chitosan polymers observed in a microsecond long molecular simulation. The solvent box is shown as a transparent glass surface and the FOV. The polymers are represented by spheres and colored based on the atom – carbon (gray), oxygen (red), nitrogen (blue), hydrogen (white). The polymers inside the solvent box (FOV) appear as bright spheres, and fade as they diffuse into neighboring periodic images in the movie. The two chitin polymers maintain a stacked conformation, while both chitosan and chitin-chitosan copolymers make short (<5Å) and long-range (<10Å) interactions with each other.

**Supplementary Movie 5: Chitin stacking.mp4**
Interaction between two chitin polymers observed in a microsecond long molecular simulation. The solvent box is shown as a transparent glass surface and the FOV. The two chitin polymers are represented by spheres and colored based on the atom – carbon (gray), oxygen (red), nitrogen (blue), hydrogen (white). The polymers in the solvent box (FOV) appear as bright spheres, and fade as they diffuse into neighboring periodic images in the movie. We observe that the two chitin polymers interact to form a stacked conformation which is stable over the length of the simulation.
